# Sexually dimorphic role of diet and stress on behavior, energy metabolism, and the ventromedial hypothalamus

**DOI:** 10.1101/2023.11.17.567534

**Authors:** Sanutha Shetty, Samuel J. Duesman, Sanil Patel, Pacific Huyhn, Sanjana Shroff, Anika Das, Disha Chowhan, Robert Sebra, Kristin Beaumont, Cameron S. McAlpine, Prashant Rajbhandari, Abha K. Rajbhandari

## Abstract

Scientific evidence underscores the influence of biological sex on the interplay between stress and metabolic dysfunctions. However, there is limited understanding of how diet and stress jointly contribute to metabolic dysregulation in both males and females. To address this gap, our study aimed to investigate the combined effects of a high-fat diet (HFD) and repeated footshock stress on fear-related behaviors and metabolic outcomes in male and female mice. Using a robust rodent model that recapitulates key aspects of post-traumatic stress disorder (PTSD), we subjected mice to footshock stressor followed by weekly reminder footshock stressor or no stressor for 14 weeks while on either an HFD or chow diet. Our findings revealed that HFD impaired fear memory extinction in male mice that received initial stressor but not in female mice. Blood glucose levels were influenced by both diet and sex, with HFD-fed female mice displaying elevated levels that returned to baseline in the absence of stress, a pattern not observed in male mice. Male mice on HFD exhibited higher energy expenditure, while HFD-fed female mice showed a decreased respiratory exchange ratio (RER). Sex-specific alterations in pro-inflammatory markers and abundance of hematopoietic stem cells were observed in chronically stressed mice on an HFD in different peripheral tissues, indicating the manifestation of distinct comorbid disorders. Single-nuclei RNA sequencing of the ventromedial hypothalamus from stressed mice on an HFD provided insights into sex-specific glial cell activation and cell-type-specific transcriptomic changes. In conclusion, our study offers a comprehensive understanding of the intricate interactions between stress, diet, sex, and various physiological and behavioral outcomes, shedding light on a potential brain region coordinating these interactions.

## Introduction

Human and animal studies show that excessive consumption of HFD contributes to weight gain [1, 2], insulin resistance [3, 4], inflammation [5–7], and an increased risk of chronic diseases, including diabetes [8], obesity [1, 9], and heart disease [10, 11]. Understanding how trauma-like stressors interact with a high-fat diet is crucial due to the high co-occurrence of conditions like PTSD and metabolic disorders. Studies have also indicated that a HFD may intensify the body’s stress response by increasing circulating levels of stress hormones [12], such as cortisol, which can contribute to heightened stress and anxiety [13, 14]. Research exploring the impact of a HFD on cognitive flexibility in rats revealed that obese rats displayed impaired cognitive flexibility, notably assessed through a set-shifting task [15]. HFD has been associated with memory impairments in a sex-specific manner. A study by Hwang et al. [16] showed that male mice fed a HFD for 12 weeks displayed impaired learning performance in contextual fear conditioning. Furthermore, research has demonstrated that HFD disrupts synaptic plasticity, particularly long-term potentiation (LTP) in the hippocampus, observed in both mice and rats [16, 17]. These findings illustrate that HFD can detrimentally influence cognitive health, resulting in compromised stress response, cognitive function, memory deficits, and synaptic dysfunction.

It is essential to highlight that research indicates potential differences in the response to a HFD between males and females in rodents, especially concerning weight gain and body composition [18]. While most studies suggest that male mice are more susceptible than females to the effects of an HFD on weight gain, metabolic changes, and cognitive function, the mechanisms at play in female mice might involve a more intricate and complex process [16, 18]. There could also be sexually dimorphic responses to the consumption of HFD in the context of chronic and acute stress. Clinical studies have revealed that symptoms of PTSD negatively impact both eating patterns and dietary behaviors. Stress-induced eating is a coping mechanism that is seen among many individuals diagnosed with PTSD, especially women [19]. This increases the likelihood of becoming obese and developing associated comorbidities that may affect health adversely [19–21]. Nevertheless, there is limited understanding regarding how unhealthy dietary habits might render an individual more vulnerable to stress, influencing mood, fear regulation, and overall mental well-being. Hence, it is crucial to comprehend the connection between diet and stress responses, specifically in a sex-specific context.

Here, we aimed at disentangling the interactions between HFD and chronic stress in male and female mice by studying the effects of intense trauma-like footshock stressor and 14 weeks HFD feeding along with repeated reminder shocks. We provide a comprehensive effect of chronic and acute stress and diet on behavior, body weight, energy metabolism, glucose homeostasis, food intake, peripheral tissue immune cell enrichments, and hypothalamic cellular state. Understanding these complex interactions is critical in addressing the rising health concerns related to high-fat diets and chronic stress. This research provides valuable insights that can guide the development of tailored prevention and intervention strategies, considering sex-specific differences, to mitigate the negative consequences of these factors and promote overall well-being in individuals facing such challenges.

## Results

### HFD led to increased fear extinction in male but not female mice

To investigate the impact of diet on stress-induced fear behavior and energy metabolism in males and females, we conducted a systematic experimental study following a specific timeline and setup (Fig. 1A). When mice (males and females) were 10 weeks old, they went through stress enhanced fear learning (SEFL) paradigm as previously described [22]. Briefly, on day 1, the animals were placed in context A and received 10 randomized footshocks. On day 2, animals were placed in context B where they received a reminder footshock at 4 minutes. After the SEFL test on day 3, the mice were divided into two groups: i) high-fat diet (HFD; 60 Kcal% fat) and ii) Chow diet (3 Kcal% fat) for 14 weeks. During this period, half of the animals in each group received a reminder shock (RS) in Context B of SEFL, serving as a repeated chronic stressor, while the remaining mice were placed in the Context B chambers without any shocks (No reminder shock; NRS) to allow for fear memory extinction. At weeks 10 and 14 of the HFD regimen, we conducted a glucose tolerance tests (GTT), administering a bolus of glucose to 4 hours fasted mice and measured blood glucose levels over two hours. At week 14, we also conducted insulin tolerance tests (ITT) to measure insulin sensitivity in response to an insulin bolus. Additionally, we placed the mice in metabolic chambers for 72 hours, as described in the methods, to perform indirect calorimetric measurements for measuring metabolic parameters in both the RS and NRS group. Subsequently, we assayed anxiety-like behaviors in the same mice by placing them in an open field-light gradient behavioral task. Finally, the animals were euthanized, and tissues (brain, brown adipose tissue (BAT), gonadal white adipose tissue (gWAT), inguinal white adipose tissue (iWAT), liver, aorta, heart, bone marrow and blood) were harvested for analysis (Fig. 1A).

**Figure 1.**
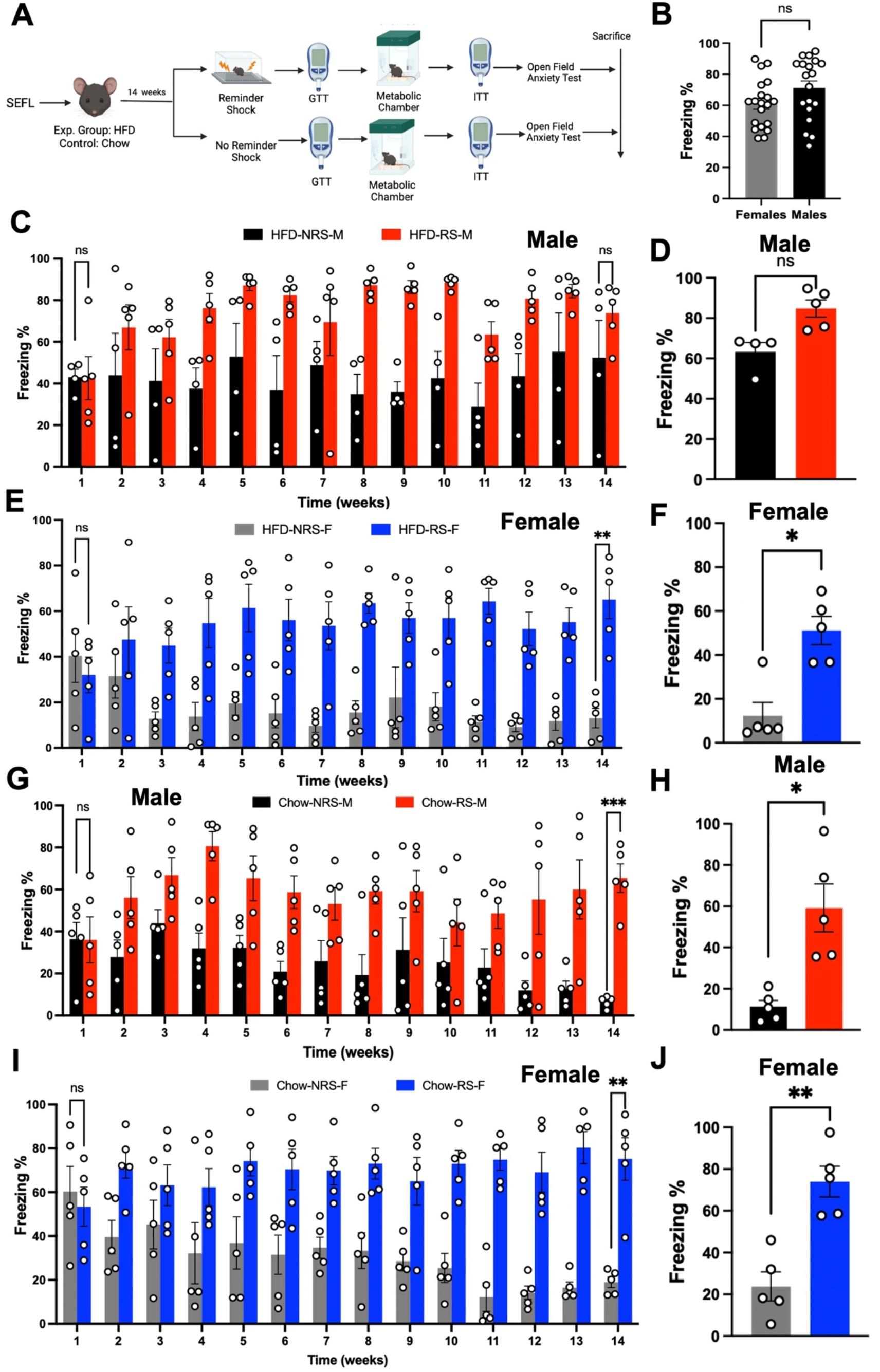
High-fat diet inhibits fear extinction in male mice. **A.** Timeline of experiment conducted to study effect of diet and repeated stress in mice (aged 10 weeks) went through stress enhanced fear learning (SEFL). Mice were then divided into two groups: one received a high-fat diet (HFD) and the other a Chow diet, for 14 weeks. During this period, half of each group were exposed to a reminder footshock stressor (RS) in Context B of SEFL, while the rest experienced no reminder shocks or fear memory extinction (NRS). Glucose tolerance test (GTT) and insulin tolerance test (ITT) were performed at weeks 10 and 14, respectively. Metabolic changes were measured using indirect calorimetric chambers, and anxiety-like behavior was assessed using an open field-light gradient task. Finally, brain and peripheral tissues were collected for analysis after sacrificing the animals for further analysis**. B.** Percent freezing at day 3 of SEFL plotted in female and male mice (N=20/19, two-tailed unpaired t-test, t=1.814, p>0.05). **C.** Percent freezing over 14 weeks of RS/NRS treatment in HFD male mice (N=4,5, two way ANOVA, F1,98 =69.18, ****p<0.0001 (*Post hoc* comparison, N=5/4, Sidak’s multiple comparison test, p>0.05). **D.** Graph represents percent freezing from the freezing test conducted after 14 weeks in HFD male mice (N=5/4, two-tailed paired t-test, t=2.518, p>0.05). **E.** Percent freezing over 14 weeks of RS/NRS treatment in HFD female mice (N=5, two way ANOVA, F1,112 =157.1, ****p<0.0001) (*Post hoc* comparison, N=5, Sidak’s multiple comparison test, p<0.05). **F.** Graph represents percent freezing from the freezing test conducted after 14 weeks in HFD female mice (N=5, two-tailed paired t-test, t=3.514, p<0.05). **G.** Percent freezing over 14 weeks of RS/NRS treatment in chow male mice (N=5, two way ANOVA, F1,112 =89.76, ****p<0.0001) (*Post hoc* comparison N=5, Sidak’s multiple comparison test, p<0.05). **H.** Graph represents percent freezing from the freezing test conducted after 14 weeks in chow male mice (N=5, two-tailed paired t-test, t=3.586, p<0.05). **I.** Percent freezing over 14 weeks of RS/NRS treatment in chow male mice (N=5, two way ANOVA, F1,112 =161.1, ****p<0.0001) (*Post hoc* comparison, N=5, Sidak’s multiple comparison test, p<0.05). **J.** Graph represents percent freezing from the freezing test conducted after 14 weeks in chow female mice (N=5, two-tailed paired t-test, t=4.771, p<0.01).

First, we assessed the percentage of freezing behavior of female and male mice before diet regimen. Our study revealed no statistically significant differences between female and male mice in percent freezing from initial SEFL paradigm exposure before diet regimen was administered with RS/NRS (Fig. 1B) (N=20,19, two-tailed unpaired t-test, t=1.814, p>0.05). However, we found a main stress effect in HFD-fed male mice male after SEFL and during the 14 weeks of diet regimen and chronic repeated stress presentation (N=4,5, two way ANOVA, F_1,98_ =69.18, p<0.05). *Post hoc* comparison reveals that by week 14 there is no difference in freezing between HFD-NRS and HFD-RS male mice (N=4,5, Sidak’s multiple comparison test, p>0.05) (Fig. 1C). This indicates that HFD-fed male mice did not show fear extinction by week 14, in the absence of footshocks as indexed by persistently high % freezing compared to HFD-RS male mice during the freezing test conducted after 14 weeks (Fig. 1D) (N=5-4, two-tailed paired t-test, t=2.518, p>0.05). Two way ANOVA on HFD-fed female mice revealed a main stress effect (N=5, two way ANOVA, F_1,112_ =157.1, p<0.05). *Post hoc* comparison shows significant reduction in freezing levels in the NRS group compared to RS group by week 14 (Fig. 1E) (N=5, Sidak’s multiple comparison test, p<0.05). Female mice also showed significant reduction in %freezing during freezing test conducted at week 14 (Fig. 1F) (N=5, two-tailed paired t-test, t=3.514, p<0.05). In the chow fed male mice groups, two way ANOVA revealed a significant stress effect (N=5, two way ANOVA, F_1,112_ =89.76, p<0.05) (Fig. 1G). *Post hoc* comparison further confirmed reduction in %freezing in chow-NRS male mice compared to chow-RS male mice (Fig. 1G) (N=5, Sidak’s multiple comparison test, p<0.05), and showed significantly decreased % freezing during freezing test done at week 14 (Fig. 1H) (N=5, two-tailed paired t-test, t=3.586, p<0.05). Similarly, we see a main stress effect in females on chow diet (N=5, two way ANOVA, F_1,112_ =161.1, p<0.05). Furthermore, *post hoc* comparisons revealed that female mice on the Chow diet also exhibited fear extinction, as indicated by decreased % freezing compared to the RS females on the Chow diet by week 14 (Fig. 1I) (N=5, Sidak’s multiple comparison test, p<0.05) and subsequently decreased %freezing in fear test (Fig. 1J) (N=5, two-tailed paired t-test, t=4.771, p<0.05). Overall, these results indicate a sex-dependent alteration in fear extinction associated with diet. In open field light gradient task there were no main effects of diet or sex in any of the HFD/chow groups (N=4,5, two way ANOVA, p>0.05) (Supplemental Fig. 1A-1E). However, there was a trend toward reduced locomotor activity during the light phase of the open field task in HFD-RS females in comparison to Chow-RS females (N=5, two way ANOVA, F_1,24_ =2.88, p=0.09) (Supplemental Fig. 1F). We did not find any such effects in male mice that received RS in either Chow/HFD-fed groups (N=5, two way ANOVA, F_1,24_ =0.054, p>0.05) (Supplemental Fig. 1G).

### HFD induced weight gain and blood glucose homeostasis in chronic stressed and non-stressed male and female mice

As shown in Fig. 2A, male and female mice on Chow diet displayed no significant effect of sex on % weight gain (N=5, Two-way ANOVA, F _3,240_ =5.698, p>0.05). However, HFD fed mice showed an effect of sex (N=4,5, Two-way ANOVA, F _32,235_ =37.12, p<0.05) (Fig. 2B). *Post hoc* comparison found a statistically significant difference in weight gain percentage between male and female mice but no effect of stress starting from week 7 (N=4,5, Tukey’s multiple comparison test, p<0.05). Next, we examined if repeated stress and HFD influenced glucose tolerance and insulin sensitivity assessed through glucose tolerance test (GTT) and insulin tolerance test (ITT), respectively. Our results showed a main effect of diet on glucose levels, with significantly enhanced glucose in female mice on HFD compared to mice on Chow diet (Fig. 2C) (N=5, Two-way ANOVA, F _3,96_ =31.53, p<0.05). *Post hoc* analysis showed that the diet-induced differences were significant at 30- to 120-minutes time points in NRS-HFD versus RS-HFD (N=5, Tukey’s multiple comparison test, p<0.05). Further, area under the curve (AUC) graph showed higher blood glucose levels in HFD-fed female mice compared to chow fed female mice without any stress effects. (Fig. 2D) (N=5, one-way ANOVA, F _3,16_ =1.041, p<0.05). By week 14, we also see a significant diet effect (N=5, two way ANOVA, F _3,96_ =13.13, p<0.05) . *Post hoc* analysis confirmed a significantly higher blood glucose level in female mice in HFD-RS group at the 30 to 120 minutes time points in comparison to the other groups (N=5, Tukey’s multiple comparison tests, p<0.05). Thereafter, blood glucose levels in female mice on HFD-NRS group returned to chow control levels while HFD-RS female mice showed sustained high blood glucose levels This was reconfirmed by significantly higher AUC in HFD-RS female mice compared to chow-fed NRS and RS female mice (N=5, one way ANOVA, F _3,16_ =0.5207, p<0.05) (Fig. 2C and 2D). We then tested blood glucose levels at 10 and 14 weeks in male mice on HFD and Chow from both NRS and RS groups. In week 10, we see a distinct diet effect on blood glucose levels with no significant effect of the stressor on blood glucose levels (Fig. 2E) (N=4,5, Two-way ANOVA, F _3,90_ =49.41, p<0.05). *Post hoc* analysis showed that the differences started to show at the 30-minute time and were sustained until 120^th^ minute of measurement (N=4,5, Tukey’s multiple comparison tests, p<0.05). AUC graph reconfirmed that HFD-fed male mice had higher blood glucose levels than chow-fed male mice (N=4,5, one way ANOVA, F _3,15_ =0.7815, p<0.05). In week 14, GTT test in male mice revealed a significant diet effect (Fig. 2F) (N=4/5, Two-way ANOVA, F _3,90_ =59.52, p<0.05). *Post hoc* analysis confirmed this by showing a sustained increase in blood glucose levels from time point 30 minutes to the 120^th^ minute (N=4/5, Tukey’s multiple comparison tests, p<0.05). Unlike in female mice on HFD, where we see with time blood glucose levels returning to baseline levels in the absence of stressors by week 14, however, we do not see that effect in male mice on HFD. AUC graph shows HFD-fed male mice had higher blood glucose levels (N=4,5, one way ANOVA, F _3,15_ =0.8658, p<0.05). We conducted ITT on 14-week HFD and chow-fed female and male mice subjected to NRS or RS. In female mice on HFD or chow diet, there was no significant diet effect on GTT irrespective of NRS and RS conditions (Fig. 2G) (N=5, two way ANOVA, F _3,16_ =1.438, p>0.05). AUC graph further confirmed no effect of diet on blood glucose levels in response to a bolus of insulin (N=5, one way ANOVA, p>0.05). Conversely, we saw an effect of diet in male mice on HFD and chow diet (Fig. 2H) (N=4/5, Two-way ANOVA, F _3,66_ =31.38, p<0.05). *Post hoc* analysis showed that male mice on HFD exhibited paradoxically low blood glucose and high insulin sensitivity compared to chow-fed mice at all time points of measurements except at 30^th^ minute (N=4/5, Tukey’s multiple comparison tests, p<0.05). AUC graph further confirms these diet differences (N=5, one way ANOVA, F _3,15_ =0.22, p<0.05). Despite the absence of distinctions between NRS and RS under HFD in male subjects, a significant reduction in insulin sensitivity was evident in Chow-RS mice compared to Chow-NRS mice. This finding indicates a potential development of insulin insensitivity due to chronic stress in male mice under a chow diet.

**Figure 2.**
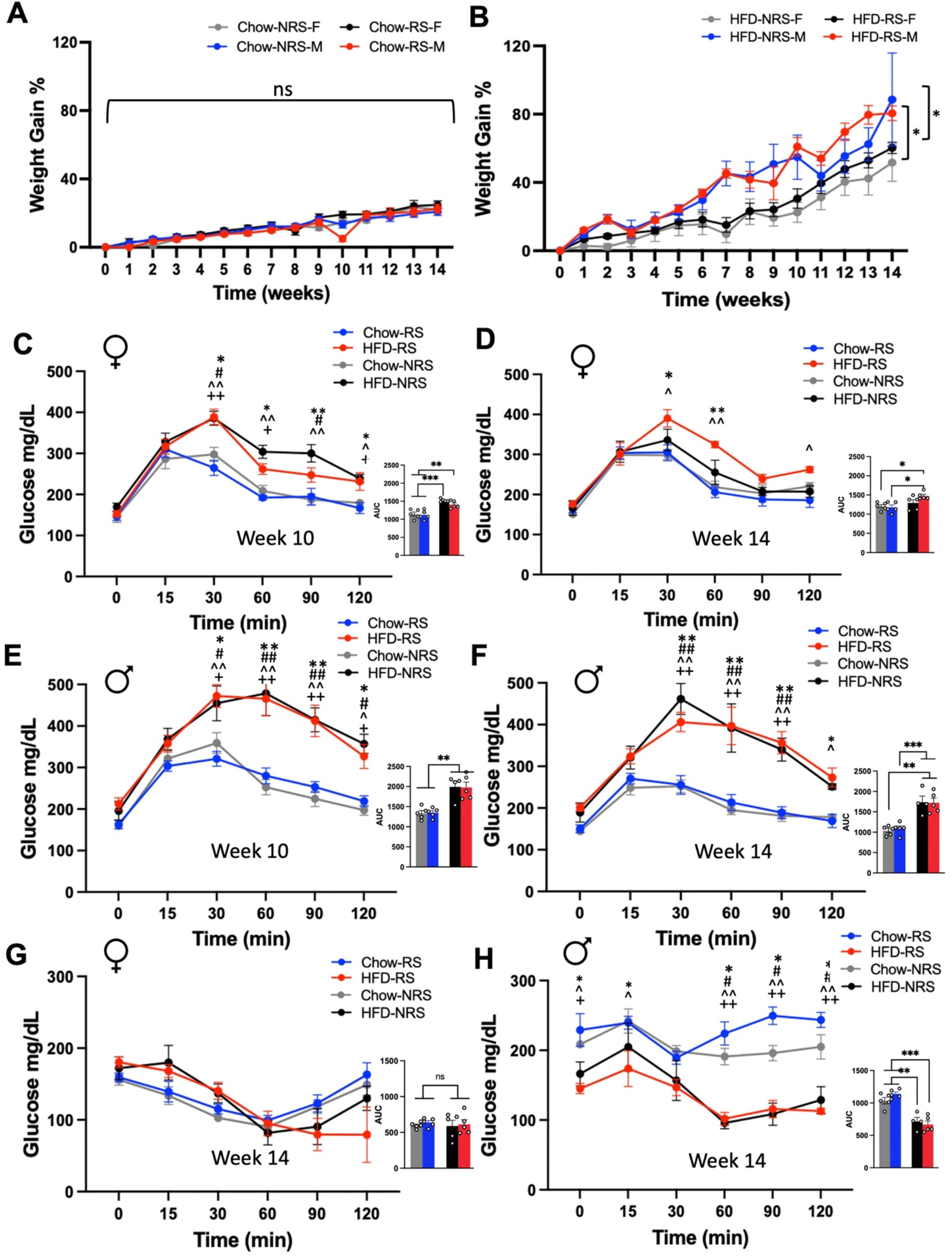
HFD induces weight gain and an increase in blood glucose in a sex-specific manner. **A.** Percent weight gain over 14 weeks plotted for chow fed mice (N=5, Two-way ANOVA, F _3,240_ =5.698, p>0.05). **B.** Percent weight gain over 14 weeks plotted for HFD fed mice (N=4/5, Two-way ANOVA, F _32,235_ =37.12, *p<0.05). **C.** Week 10 plasma glucose levels during GTT test following 4 hours of fasting in HFD/chow female mice (N=5, two way ANOVA, F _3,96_=31.53, ****p<0.0001) (*Post-hoc* comparison, N=5, Tukey’s multiple comparison test, p<0.05). AUC graph from 10 week GTT test in HFD/chow fed female mice (N=5, one way ANOVA, F _3,16_ =1.041, ****p<0.0001) (*Post hoc* comparison, N=5, Tukey’s multiple comparison test, p<0.05) **D.** Week 14 plasma glucose levels during GTT test following 4 hours of fasting in HFD/chow female mice (N=5, two way ANOVA, F _3,96_=13.13, ****p<0.0001) (*Post hoc* comparison, N=5, Tukey’s multiple comparison test, p<0.05). AUC graph from 14 week GTT test in HFD/chow female mice (N=5, one way ANOVA, F _3,16_=0.527, *p<0.05) (*Post hoc* comparison, N=5, Tukey’s multiple comparison test, **p<0.01, ***p<0.001). **E.** Week 10 plasma glucose levels during GTT test following 4 hours of fasting in HFD/chow male mice (N=5, two way ANOVA, F _3,90_=49.41, ****p<0.0001) (*Post hoc* comparison, N=4/5, Tukey’s multiple comparison tests, p<0.05). AUC graph from 10 week GTT test in HFD/chow fed male mice (N=5, one way ANOVA, F _3,15_=0.7185, ***p<0.001) (*Post hoc* comparison, N=4/5, Tukey’s multiple comparison test, **p<0.01). **F.** Week 14 plasma glucose levels during GTT test following 4 hours of fasting in HFD/chow male mice (N=5, two way ANOVA, F _3,66_=55.89, ****p<0.0001) (*Post hoc* comparison, N=4/5, Tukey’s multiple comparison tests, p<0.05). AUC graph from 14 week GTT test in HFD/chow fed male mice (N=5, one way ANOVA, F _3,15_=0.8658, ****p<0.0001) (*Post hoc* comparison, N=4/5, Tukey’s multiple comparison test, **p<0.01, ***p<0.001). **G.** Week 14 plasma glucose levels during ITT test following 4 hours of fasting in HFD/chow female mice (N=5, Two-way ANOVA, F_3,93_ = 0.5491, p>0.05). AUC graphs from 14 week ITT test in HFD/chow fed female mice (N=5, one way ANOVA, F _1,16_=1.438, p>0.05) **H.** Week 14 plasma glucose levels during ITT test following 4 hours of fasting in HFD/chow male mice (N=5, two way ANOVA, F _3,66_=31.38, ****p<0.0001) (*Post hoc* comparison, N=4/5, Tukey’s multiple comparison tests, p<0.05). AUC graphs from 14 week ITT test in HFD/chow fed male mice (N=5, one way ANOVA, F _3,15_=0.22, ****p<0.0001) (*Post hoc* comparison, N=5, Tukey’s multiple comparison test, **p<0.01, ***p<0.001). (*Compares Chow-No repeated Shock to HFD-repeated shock, #Compares Chow-No repeated shock to HFD-No repeated shock, ^Compares Chow-Repeated Shock to HFD-Repeated Shock, +Compares Chow-Repeated Shock to HFD-No repeated Shock).

### Differential effects of stress and HFD on energy balance

In addition to glucose and insulin homeostasis, stress and HFD are also known to modulate energy homeostasis, potentially leading to obesity and associated metabolic disorders. To investigate the effects of diet and repeated shocks on energy metabolism, we placed the chow-fed and HFD-fed RS and RS mice in metabolic chambers for 72 hours to measure indirect calorimetric parameters such as energy expenditure (EE), respiratory exchange ratio (RER), food intake, and locomotion. We omitted the initial 12 hours of acclimation data. We saw a main effect of sex in the energy expenditure of HFD fed mice (Fig. 3A) (N=4,5, one way ANOVA, p<0.05). *Post hoc* comparison found that under HFD, male mice exhibited significantly higher EE than females, but EE was comparable between RS and non-repeated shocks NRS in both males and females signifying no stress effects (N=4,5, Tukey’s multiple comparison test, p<0.05). However, there is no sex or stress effect on overall EE, as it was comparable between males and females under a chow diet, with a tendency towards an increase in EE under RS for both sexes (Fig. 3B) (N=5, One way ANOVA, p>0.05). We then examined fuel utilization in males and females under different diets and stress levels by measuring RER levels. RER, signifies the ratio between the body’s production of carbon dioxide (CO2) and the absorption of oxygen (O2), serves as an indicator of the type of energy source (such as carbohydrates or fats) metabolized to provide energy to the body. We saw a main effect of sex in RER in HFD fed mice (Fig. 3C) (N=4/5, One way ANOVA, p<0.05), with *post hoc* comparisons we found marked differences in RER values were observed between male mice and female mice, with female mice under both RS and NRS showing significantly lower (N=4,5, Tukey’s multiple comparison test, p<0.05). On chow diet, we find no sex or stress effects on RER values suggesting that they were comparable between male mice and female mice (Fig. 3D) (N=5, One way ANOVA, p>0.05). Despite higher EE and RER values in males, no main effect of sex or stress was found in total food consumption between male mice and female mice fed HFD (Fig. 3E) (N=4/5, One way ANOVA, p>0.05). We did not see any sex or stress effects in total food consumed in female or male mice on chow diet (Fig. 3F) (N=5, One way ANOVA, p>0.05). We also measured the locomotor activity of male and female mice by calculating pedestrian locomotion inside the cages. Interestingly, we saw a main effect of sex in locomotion in HFD fed mice (Fig. 3G) (N=5, One way ANOVA, p<0.05), with *post hoc* comparisons we found that RS mice exhibited lower locomotion activity compared to NRS under HFD, and NRS and RS female mice showed comparable locomotion (N=4,5, Tukey’s multiple comparison test, p<0.05). We also saw a main sex effect in pedestrian locomotion in chow fed mice (Fig. 3H) (N=5, One way ANOVA, p<0.05). *Post hoc* analysis revealed that both female mice groups exhibited higher locomotion than male mice, with a significant difference between RS male mice and female mice (N=4,5, Tukey’s multiple comparison test, p<0.05).

**Figure 3.**
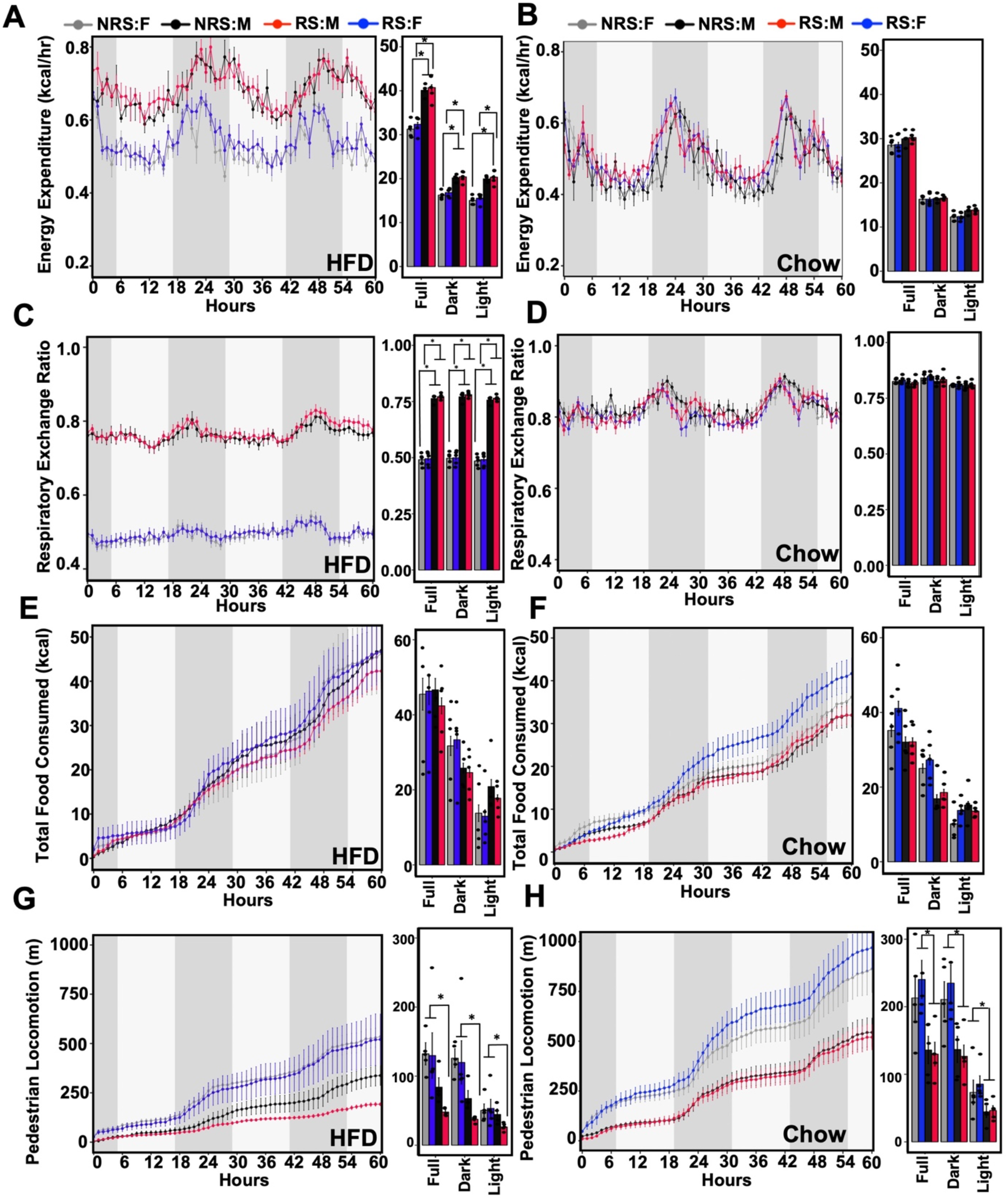
HFD and repeated shock causes a sexually dimorphic disruption in metabolism. **A.** Energy expenditure (kCal/hr) of HFD fed mice over 60 hours in the metabolic chamber (N=4,5, one way ANOVA, p<0.05) (*Post hoc* comparison, N=4,5, Tukey’s multiple comparison test, *p<0.0**5**). **B.** Energy expenditure (kCal/hr) of chow fed mice over 60 hours in the metabolic chamber (N=5, One way ANOVA, p>0.05). **C.** Respiratory exchange ratio (RER) of HFD fed mice over 60 hours in the metabolic chamber (N=4/5, One way ANOVA, p<0.05) (*Post hoc* comparisons, N=4,5, Tukey’s multiple comparison test, *p<0.05). **D.** RER of chow fed mice over 60 hours in the metabolic chamber (N=5, One way ANOVA, p>0.05). **E.** Total food consumption in HFD fed mice over 60 hours in the metabolic chamber (N=4/5, One way ANOVA, p>0.05). **F.** Total food consumption in chow fed mice over 60 hours in the metabolic chamber (N=5, One way ANOVA, p>0.05). **G.** Pedestrian locomotion in HFD fed mice over 60 hours in the metabolic chamber (N=5, One way ANOVA, p<0.05) (*Post hoc* comparisons, N=4,5, Tukey’s multiple comparison test, *p<0.05). **H.** Pedestrian locomotion in chow fed mice over 60 hours in the metabolic chamber (N=5, One way ANOVA, p<0.05), (*Post hoc* comparison, N=4,5, Tukey’s multiple comparison test, *p<0.05).

### Combined effect of HFD and acute stress on behavior, glucose homeostasis, and energy metabolism

To investigate the impact of HFD on acute stress-induced fear-related behaviors and energy metabolism, we conducted a comprehensive experimental study following a specific timeline and setup (Fig. 4A). At 10 weeks of age, we segregated the mice into the High Fat Diet (HFD) or Chow groups for 10 weeks. Following the 10-week dietary intervention, the mice were exposed to either an acute or no-stress paradigm, as outlined in our experimental methods. Additionally, following the procedures detailed in the methods section, we utilized metabolic chambers to conduct indirect calorimetric measurements to evaluate the metabolic processes. Subsequently, GTT tests were performed on those mice. As shown in Fig. 4B, there was both a sex effect and a diet effect on weight gain percentage from baseline (N=10, two way ANOVA, F_3,528_ =188.3 & F_10,528_ =54.75, p<0.05). As expected, HFD fed animals gained more weight than chow fed control animals and these differences started at week 2 of diet regimen (N=10, Tukey’s multiple comparison test, p<0.05). However, males in both diet groups gained weight at a significantly higher rate than female counterparts from the same group (N=10, Tukey’s multiple comparison test, p<0.05). Next, we looked at fat mass in these animals, and two-way ANOVA showed main diet effect (Fig. 4C) (N=10, Two-way ANOVA, F_1,48_ = 17.69, p<0.05). *Post hoc* analysis found that only males on HFD showed a significantly higher percentage of fat mass compared to males on the Chow diet (N=10, Tukey’s multiple comparison test, p<0.05). Despite higher weight gain in females on HFD, they did not show a difference in fat mass compared to females on the chow diet (N=10, Tukey’s multiple comparison tests, p>0.05). The analysis of freezing behavior in these animals revealed a main effect of shock but no main effect of sex or diet (Fig. 4D) (N=5, One-way ANOVA, F_7,44_ = 1.388, p<0.05). *Post hoc* analysis revealed these differences between the Chow-NS-F v. Chow-S-F, Chow-NS-M v. Chow-S-M, HFD-NS-F v. HFD-S-F, HFD-NS-M v. HFD-S-M (N=5, Tukey’s multiple comparison test, p<0.05). GTT tests after SEFL showed a main diet effect in female mice on either S or NS conditions (Fig. 4E) (N=5, two way ANOVA, F_3,144_ =12.28, p<0.05). *Post hoc* comparison further revealed this sex effect was pronounced during the 30^th^-90^th^ minute of measurement between HFD-S female mice and chow fed mice in both groups (N=5, Tukey’s multiple comparison test, p<0.05). AUC graph also reveals a significantly higher blood glucose levels in HFD-S female mice compared to chow-S female mice (N=5, one way ANOVA, F_3,16_ =0.1784, p<0.05). We also saw a main effect of diet in male mice (Fig. 4F) (N=5, two way ANOVA, F_3,120_ =9.057, p<0.05). *Post hoc* comparison revealed this difference in HFD fed male mice and chow fed male mice was exhibited from 30-90 minutes of measurement (N=5, Tukey’s multiple comparison test, p<0.05). AUC graph for the GTT test, also confirmed a diet effect on blood glucose levels (N=5, one way ANOVA, F_3,16_ =0.6993, p<0.05). Next, we placed male and female mice that were on either HFD or a chow diet and under S and NS conditions in metabolic chambers to measure changes in EE. We did not observe a sex effect in EE between HFD female and male mice under either the S or NS conditions (Fig. 4G) (N=5, One way ANOVA, p>0.05). We also do not observe a sex effect in EE in chow fed animals (Fig. 4H) (N=5, One way ANOVA, p>0.05). We also looked at other metabolic parameters including RER, total food consumption and locomotor activity with sex grouping (Supplemental Fig. 2). We saw an effect of diet in RER in both female and male mice under both chow and HFD condition without any stress effects (Supplemental Fig. 2A & 2B). The impact of stress on HFD-fed females is evident in the reduced food intake, whereas Chow-S female mice exhibit increased food consumption compared to their NS counterparts. (Supplemental Fig. 2C).

**Figure 4.**
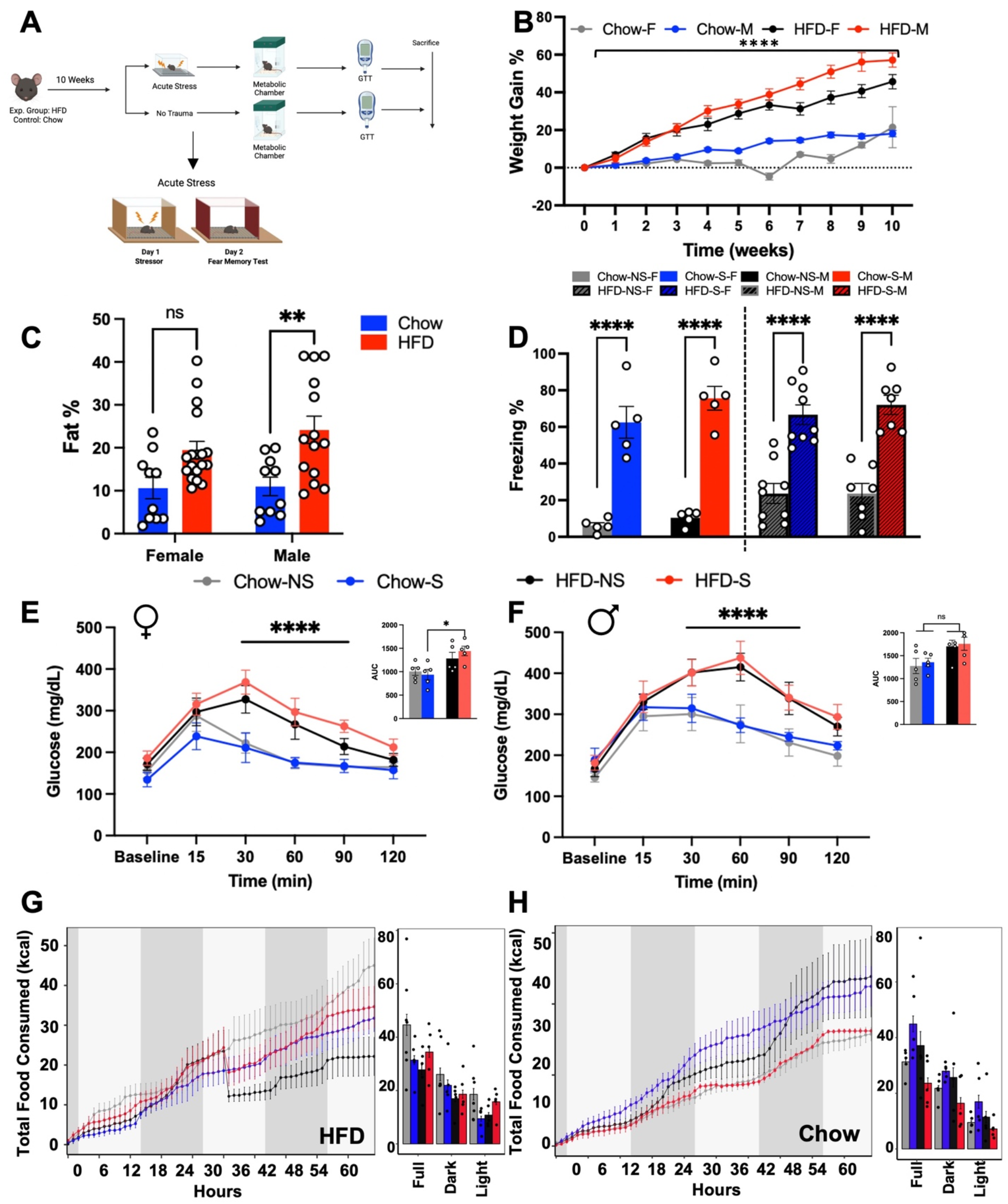
HFD has no effect on acute stressed induced fear behavior but shows sex specific changes in metabolism. **A.** Experimental design of the comprehensive 10-week study conducted to investigate the role of diet and cute stress. Mice were placed in HFD or Chow groups, followed by exposure to acute stress or no stress. Metabolic chambers were used for indirect calorimetric measurements. A GTT test was next conducted to measure blood glucose levels after 4 hours of fasting. The study concluded with animal sacrifice and tissue harvest. **B.** Weight gain percent plotted through 10 weeks of either HFD/chow diet regimen (N=10, Two way ANOVA, F3,528 = 188.3, ****p<0.0001). **C.** Percent fat mass in HFD/chow fed mice, revealing a sex difference (N=10, Two way ANOVA, F1,48 = 17.69, **p<0.01) (*Post hoc* comparison, N=10, Tukey’s multiple comparison test, **p<0.01). **D.** Analysis of freezing behavior in these animals showed a group effect between the no-shock and shock groups (N=5, One way ANOVA, F7,44 = 1.388, ****p<0.0001) (*Post hoc* comparison, N=5, Tukey’s multiple comparison test, Chow-NS-F v. Chow-S-F, Chow-NS-M v. Chow-S-M, HFD-NS-F v. HFD-S-F, HFD-NS-M v. HFD-S-M; ****p<0.0001). **E.** Plasma blood glucose levels from GTT test performed at week 10 in HFD/Chow – NS/S females (N=5, two way ANOVA, F_3,96_ = 19.55 ****p<0.0001). AUC graph for week 10 GTT test in HFD/chow fed female mice (N=5, One way ANOVA, F_3,16_ = 4.789, *p<0.05). **F.** Plasma blood glucose levels from GTT test performed at week 10 in HFD/Chow – NS/S males (N=5, two way ANOVA, F_3,96_ = 13.3, ****p<0.0001). AUC graph for week 10 GTT test in HFD/chow fed male mice (N=5, One way ANOVA, F_3,16_ = 3.248, p>0.05). **G.** Energy expenditure (EE) (kCal/hr) in HFD fed mice under either the S or NS conditions (N=5, One way ANOVA, p>0.05). **H.** Energy expenditure (EE) (kCal/hr) in chow fed mice under either the S or NS conditions (N=5, One way ANOVA, p>0.05).

### Repeated stress induces differential abundance of myeloid lineage cells in bone marrow and inflammatory cells in the blood, gWAT, aorta, and heart

To investigate the potential systemic inflammation resulting from repeated shock, we first evaluated the expression of pro-inflammatory genes in the gWAT and liver utilizing quantitative polymerase chain reaction (qPCR). Among the assessed genes, Monocyte Chemoattractant Protein-1 (*Mcp1*), a chemokine involved in regulating monocyte/macrophage migration and infiltration, was particularly interesting due to its implication in metabolic processes [38, 39]. However, we did not see any stress or sex effects on gene expression of *Mcp1* (N=9,10, two way ANOVA, p>0.05) (Supplemental Fig. 3A & 3E). Interestingly, we did see a sex and stress effect on gene expression levels of pro-inflammatory cytokine *Il-1β* (N=9,10, two way ANOVA, F_1,15_ = 9.796 &11.08 respectively, p<0.05) and cytokine *Il-12p40* (N=9,10, two way ANOVA, F_1,15_ = 9.796 & 11.08 respectively, p<0.05) in HFD-fed animals (Supplemental Fig. 3B & 3C). In our analysis on the liver, we see no effects of sex or stress on *Mcp-1, Il-1β, Il-12p40* or *TNF α* gene expression levels (N=9, two way ANOVA, p<0.05) (Supplemental Fig. 3E-H).

We next investigated leukocyte abundance in multiple peripheral tissues in RS and NRS HFD-fed mice, aiming to gain deeper insights into the impact of stress and diet on systemic inflammation. Inflammation is an important contributor to metabolic abnormalities which together can contribute to the progression of cardiovascular disease (CVD) [23]. Although we did not test CVD phenotype in this study, testing leukocyte abundance in organs relevant to CVD may inform on the relevance of our findings to this disease. To this front, we employed flow cytometric analysis (FACS) of cells collected from bone marrow (BM), blood, gWAT, aorta and heart tissues of HFD-fed RS and NRS female and male mice as detailed in the methods. Gating strategies for the flow cytometry data analysis is shown in Supplemental Fig 4A-4E. Multipotent progenitors (MPPs) in the BM, including MPP2 and MPP3, generate myeloid and lymphoid cell populations. MPP2 tends to differentiate into lymphocytes and megakaryocytes, whereas MPP3 induces differentiation into monocytes and granulocytes [24]. We found a main sex effect in the MPP2 cell population in BM of HFD fed mice (N=9, two-way ANOVA, F_1,14_ = 290.1, p<0.05) (Fig. 5A). The *post hoc* comparison showed significant reduction in MPP2 cell population in HFD-RS female mice compared to HFD-NRS male mice (N=5, Tukey’s multiple comparison test, p<0.05) (Fig. 5A). Additionally, we also observe a significant decrease in MPP2 cells in HFD-male mice in comparison to HFD-female mice irrespective of stress classification (N=4/5, Tukey’s multiple comparison test, p<0.05). We saw similar main sex effects with population of MPP3 progenitor population in BM of HFD fed mice (N=9, two-way ANOVA, F_1,14_ = 34.48, p<0.05) (Fig. 5A). While *post hoc* analysis shows decreasing trend in HFD-RS female mice in comparison to HFD-NRS female mice it was not statistically significant (N=5, Tukey’s multiple comparison test, p=0.08). A significant reduction in MPP3 cells were seen in HFD fed male mice in comparison to HFD fed female mice irrespective of stress grouping (N=4/5, Tukey’s multiple comparison test, p<0.05). Next, we looked at the downstream products of these MPPs, namely the granulocyte-monocyte precursors (GMPs) and the monocyte-dendritic progenitor cells (MDPs) in the BM and see a statistically significant sex effect (Fig. 5C and 5D). Hematopoietic stem cells, including MPP2s and MPP3s go through a series of steps as they become increasingly committed, ultimately giving rise to multipotent common myeloid progenitor cells (CMPs), which, in turn, have the potential to differentiate into GMPs and MDPs and ultimately mature leukocytes [41]. Quantification of GMPs in HFD-fed animals shows a main sex effect (N=9, two-way ANOVA, F_1,14_ = 42.52, p<0.05). *Post hoc* comparison reveals a statistically significant decrease in GMPs in HFD-fed female mice in comparison to their male counterparts (N=5-4, Tukey’s multiple comparison test, p<0.05), however, we do not see any stress effects. On the other hand, we see a main stress effect in MDPs cell population in HFD-fed groups (N=9, two-way ANOVA, F1,14 = 6.017, p<0.05). The *post hoc* comparisons show a significant reduction in MDP levels in HFD-RS males in comparison to HFD-NRS females (N=5-4, Tukey’s multiple comparison test, p<0.05) (Fig. 5C and 5D).

**Figure 5.**
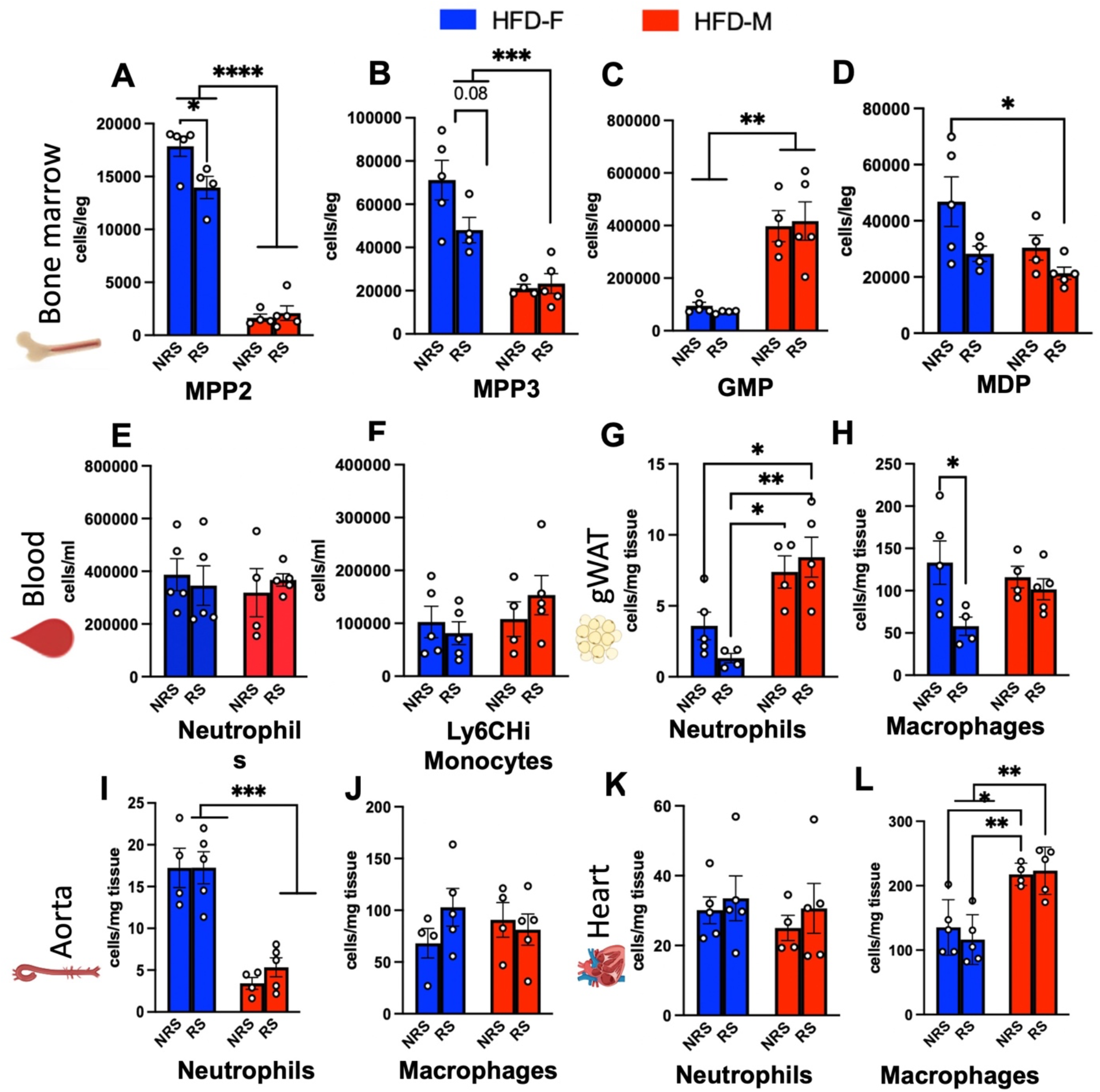
Repeated shock and HFD induces differential changes in peripheral myeloid lineage cells and inflammatory markers in a sexually dimorphic manner. **A.** In the bone marrow (BM), quantification of MPP2 progenitor population in HFD fed mcie (N=9, two-way ANOVA, F1,14 = 290.1, ****p<0.0001) (*Post hoc* comparison, N=5, Tukey’s multiple comparison test, *p<0.05 & ****p<0.0001). **B.** Quantification of MPP3 progenitor population in BM of HFD fed mice (N=9, two-way ANOVA, F1,14 = 34.48, ****p<0.0001) (*Post hoc* comparison, N=5, Tukey’s multiple comparison test, p=0.08 & ***p<0.001). **C.** Quantification of granulocyte monocyte progenitor (GMPs) cells in HFD fed mice (N=9, two-way ANOVA, F1,14 = 42.52, ****p<0.0001) (*Post hoc* comparison, N=4/5, Tukey’s multiple comparison test, **p<0.01). **D.** Quantification of monocyte-dendritic cell progenitors (MDPs) in BM in HFD fed mice (N=9, two-way ANOVA, F1,14 = 6.017, *p<0.05) (*Post hoc* comparison, N=5, Tukey’s multiple comparison test, *p<0.05). **E.** Quantification of neutrophils in the blood of HFD mice (N=9, two-way ANOVA, p>0.05). **F.** Quantification of Ly6Chi Monocytes in the blood of HFD fed mice (N=9, two-way ANOVA, p>0.05). **G.** Quantification of neutrophils in the gonadal white adipose tissue (gWAT) of HFD fed mice (N=9, two-way ANOVA, F1,14 = 24.65, ***p<0.001) (*Post hoc* comparison, N=5, Tukey’s multiple comparison test, *p<0.05 & **p<0.01). **H.** Quantification of macrophages in the gWAT of HFD fed mice (N=9, two-way ANOVA, F1,14 = 6.296, *p<0.05) (*Post hoc* comparison, N=4/5, Tukey’s multiple comparison test, *p<0.05). **I.** Quantification of aortic neutrophils of HFD fed mice (N=9, two-way ANOVA, F1,14 = 59.59, ****p<0.0001) (*Post hoc* comparison, N=4/5, Tukey’s multiple comparison test, ***p<0.001). **J.** Quantification of macrophages in the aorta of HFD mice (N=4/5, Tukey’s multiple comparison test, p>0.05). **K.** Quantification of neutrophils in the heart of HFD mice (N=9, two-way ANOVA, p>0.05). **L.** Quantification of macrophage in the heart of HFD fed mice (N=9, two-way ANOVA, F1,14 = 32.04, ****p<0.0001) (*Post hoc* comparison, N=4/5, Tukey’s multiple comparison test, *p<0.05 & **p<0.01).

For a more comprehensive understanding of peripheral alterations in response to both stress and diet, we analyzed blood samples obtained from these respective groups. We do not see any effect of sex or stress on neutrophils or Ly6Chi monocyte cell population in blood samples (Fig. 5E and 5F) (N=5-4, One way ANOVA, p>0.05). Next, we analyzed neutrophils and macrophage cell populations in gWAT, aorta and heart tissue (Fig. 5G-5L), critical to systemic metabolic regulation. A significant sex effect was seen in gWAT neutrophil levels (Fig. 5G) (N=9, One way ANOVA, F_1,14_=24.65, p<0.05) but a stress effect in macrophage cell population (Fig. 5H) (N=9, One way ANOVA, F_1,14_=6.296, p<0.05). In *post hoc* comparisons we saw an increase in neutrophil levels in HFD-fed male mice in comparison to HFD-RS female mice in both stress groups (N=5-4, Tukey’s multiple comparison test, p<0.05) and an increase in neutrophil levels in HFD-NRS female mice in comparison to HFD-RS male mice (N=5, Tukey’s multiple comparison test, p<0.05). While in the aorta we saw a converse sex effect on neutrophil levels (N=9, two-way ANOVA, F1,14 = 24.65, p<0.05) (Fig. 5I) with HFD-female showing a reduction in neutrophil levels in comparison to HFD-fed males (N=9, two way ANOVA, F_1,14_=59.59, p<0.05). The *post hoc* analysis shows a significantly higher levels of neutrophils in HFD-fed female mice than male mice irrespective of stress grouping (N=5, Tukey’s multiple comparison test, p<0.05). We did not see any sex, or stress effects in macrophages levels in aorta (Fig. 5J) (N=5-4, One way ANOVA, F_3,14_=0.2060, p>0.05). In the heart, we did not see sex or stress effects in neutrophil levels in HFD animals (Fig. 5K) (N=9, two way ANOVA, F_1,14_=0.3649, p>0.05). However, we saw a significant sex effect in macrophage levels in the heart (Fig. 5L) (N=9, two way ANOVA, F_1,14_=32.04, p<0.05). *Post hoc* analysis showed that HFD-NRS females had significantly lower levels of macrophages than HFD-fed male mice (N=5-4, Tukey’s multiple comparison test, p<0.05) and HFD-RS females hadsignificantly lower levels of macrophages than HFD-fed male mice (N=5-4, Tukey’s multiple comparison test, p<0.05).

We also analyzed the tissues from the chow-fed animals, and our observations revealed minimal sex or stress impacts on the myeloid cell lineage population and pro-inflammatory markers (Supplemental Fig. 4F-4M). Interestingly, we noted a main sex effect in the population of GMPs and MDPs cells in chow-fed animals (N=10, two way ANOVA, F_1,16_ = 9.504 and 5.604 respectively, **p<0.01 & *p<0.05 respectively). *Post hoc* comparison teased out differences in chow-NRS males in comparison to chow-NRS females (N=5, Tukey’s multiple comparison test,* p<0.05) (Supplemental Fig. 4C and 4D). These findings provide additional support for the detrimental influence that high-fat diet, when combined with chronic stress, can have on these biological parameters.

### VMH snRNA-seq shows differential cell states under chronic repeated stress and HFD in male and female mice

Given discernible changes in behavior and energy metabolism due to dietary and stress-related challenges from our results, we focused on analyzing the ventromedial hypothalamus (VMH). VMH is known to coordinate glucose and energy homeostasis in response to metabolic need in a sexually dimorphic manner [25] [26, 27]. It is also known to mediate fear behavior in rodents [29, 30]. Therefore, we proceeded to examine the role of VMH for studying meaningful cell state and cell-to-cell gene expression variability. We conducted snRNA-seq to examine the impact of repeated shock and a HFD on the VMH regions of both male and female mice (Fig. 6A). Nuclei were isolated from the VMH regions of HFD-RS male and female mice (pooled from 5 mice each). The single nuclei suspension (n ∼ 10,000) underwent snRNA-Seq using the 10XGenomics platform, and libraries were sequenced with a dedicated 1 billion reads per sample. We utilized the 10X Genomics data processing platform and SeuratV3 to generate cell clusters and identities (refer to Materials and methods) (Fig. 6A). To classify VMH populations based on gene expression, we conducted cluster analysis, as illustrated by Uniform Manifold Approximation and Projection (UMAP) plots (Fig. 6B). These plots enabled us to distinguish distinct clusters of GABAergic neurons, mature neurons, dopaminergic neurons, oligodendrocyte precursors, astrocyte populations, glutamergic neurons, oligodendrocyte populations, endothelial cells, and microglia. Notably, based on marker genes, we identified two populations of astrocytes and oligodendrocytes (Fig. 6B). Further examination through UMAP and gene network plots revealed that each cluster uniquely expressed marker genes, demonstrating a preferential expression in individual clusters (Sup. Fig. 5A & 5B). To gain insight into sex differences in VMH remodeling, we segregated the cumulative UMAP plot into female and male mice (Fig. 6C). The UMAP indicated overall comparable qualitative changes in the relative proportions of VMH clusters between male and female nuclei (Fig. 6C). To characterize the differences in cell fraction between males and females, we calculated cluster percentages in relation to the overall combined dataset. We observed variations in astrocytes, oligodendrocytes, microglial cells, GABAergic neurons, and mature neurons (Fig. 6D). Transcriptional analyses based on gene signatures showed an active transcriptional network involving transcription factors such as NFIB, MEIS1, SOX10, and FOXO1 in glial cells. This suggests a significant impact of diet and chronic stress on the transcriptional program of glial cells (Fig. 6E). Further analysis involved subsetting glial cells, including microglia, astrocytes, and oligodendrocytes, revealing higher percentages of microglia and oligodendrocytes in females, marked by *Ctss* and *Mag*, and an astrocyte population in males. Interestingly, neuropeptide *Pomc* and astrocyte marker *Gfap* were notably higher in male astrocytic populations (Fig. 6F). To delineate transcriptional differences in glial cell populations between males and females, we performed differential gene expression analysis on oligodendrocytes, microglia, and astrocyte populations. The volcano plot in Fig. 6G illustrates highly upregulated genes in the microglia of female mice, including *Xist, Hcrt,* and *Ctnap5a*, compared to males. For the astrocytic population, highly significant genes in males were *Nrg3, Lingo2,* and *Grid2* (Fig. 6H) . In female mice, the oligodendrocyte population consisted of genes such as *Pmch, Tsix,* and *Rplp1* (Fig. 6I). To correlate gene expression profiles with pathway analysis, we performed gene ontology for differentially expressed genes in the overall neuronal population, astrocytes, and microglia, correlating with cell type activation pathways. Overall neuroinflammation was comparable between males and females (Fig. 6J), but males showed an increase in *App* and *Ldlr* expression, while females showed an increase in *Lrp1* and *Mapt* expression, indicative of differential activation of astrocytes lipid metabolism (Fig. 6K). Consistent with previous results, female mice showed overall activation of microglia, with increased expression of *Nr1d1, Sty11,* and *Clu* (Fig. 6L). Along these observations, examination of cell cluster expression association with genome wide association studies (GWAS) traits using program MAGMA [42] showed a higher correlation of female with Alzheimer frequency than males under chronic stress and HFD (Sup. Fig. 5C). To further study how neuronal and non-neuronal cells interact in male and female mice under chronic stress and HFD, we performed cell-cell communication (ligand-receptor interaction) analysis using CellChat [28]. We found markedly more active communication between cell types in females than males (Fig. 6M and Sup. Fig. 5D). Females exhibited strong mature neuron autocrine effects, with robust interactions between mature neurons, glutamergic neurons, and tanycytes (Fig. 6M). Oligodendrocyte precursor connections were comparable between males and females, while in females, oligodendrocytes showed stronger connections with glutamergic neurons, tanycytes, dopaminergic neurons, and astrocytes (Fig. 6M). Female mice also showed stronger connections from microglia to glutamergic neurons, oligodendrocyte precursors, and astrocytes (Fig. 6M). The connections from tanycytes were similar between males and females, while dopaminergic neuronal connections with oligodendrocyte precursors were stronger in males (Fig. 6M). We further analyzed highly expressed ligands from our dataset in relation to their interaction with various cell types, plotted as a heatmap (Fig. 6N). Sex-specific differences in cellular responses to ligands were observed, with factors such as *Negr* and *Epha* showing stronger interactions with dopaminergic neurons, *Sema6* with dopaminergic and glutamergic neurons, and laminin with astrocytes in females (Fig. 6N). In males, the intensity of ligand and cellular connections was overall more pronounced, with *Cntn* to dopaminergic neurons, *Epha* to mature neurons, *Cdh* to glutamergic neurons, *Ptprm* to astrocytes, oligodendrocytes, mature neurons, and microglia, *Sema6* to oligodendrocyte precursors, and *Bmp* to endothelial cells (Fig. 6N).

**Figure 6.**
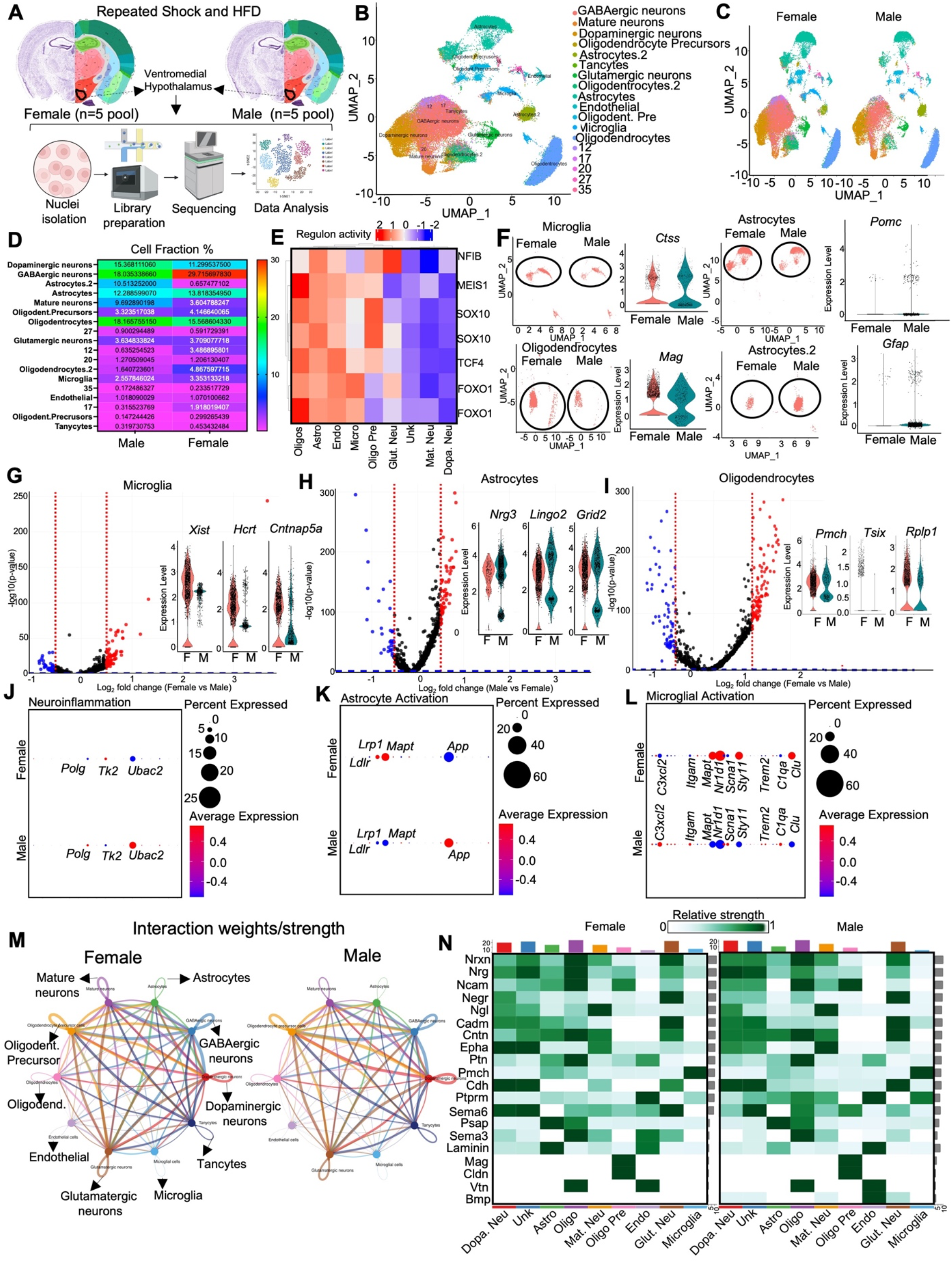
Repeated shock and HFD induces differential changes in the VMH of male and female mice. **A.** Illustration outlining the snRNA-seq procedure utilizing the 10XGenomics platform workflow, which involves isolating nuclei from the VMH of RS:HFD male and female mice (N=5 pooled). **B.** UMAP plot displaying single cells from this study, color-coded by cell type, with cell types identified based on the expression of canonical marker genes. **C.** UMAP plot of single cells from the VMH, color-coded by cell type and segregated by sample. **D.** Heatmap depicting the relative fractions of each cell type in each sample. **E.** Heatmap illustrating the regulon activity of indicated transcription factors, indicating the intensity of gene regulation in specific cell types from the VMH of males and females. **F.** UMAP plot of indicated cell types from the VMH, separated by sample, accompanied by adjacent violin plots displaying expression levels of specified genes. **G,H & I.** Volcano plot illustrating differentially expressed genes (DEGs) with fold change plotted against p-value, comparing female vs male in microglia (G), male vs female in astrocytes (H), and female vs male in oligodendrocytes (I). Violin plots highlight highly upregulated genes. **J, K & L.** Pathway analysis derived from DEGs, showing neuroinflammation (J), astrocyte activation (K), and microglial activation (L). **M.** Interactome representation based on DEGs from male and female mice, indicating the strength of interaction between specified cell types. **N.** Heatmap exhibiting indicated ligand-cell interactions in male and female mice.

In conclusion, our snRNA-seq data, for the first time, revealed the influence of chronic stress and a high-fat diet on the VMH in male and female mice. While the sex differences were not overtly pronounced, there were major changes in the transcriptomic state of glial cells between males and females that could regulate neuronal function to influence behavior and energy metabolism.

## Discussion

Our studies show a previously unexplored effect of diet and chronic stress on fear-related behavior and various metabolic, immune, and hypothalamic implications. We find that male mice on HFD show inhibition in fear extinction. While it is not surprising that HFD fed mice show higher than expected blood glucose levels in GTT test, we see that this is reversible in female mice on HFD in the absence of the stressor. Further on the metabolic front, we found that female mice on HFD have lower RER indicating sourcing of energy from fat instead of carbohydrates. We also see higher EE in HFD fed male mice in comparison to HFD fed female mice. Chow diet had minimal impact on the range of metabolic parameters tested, however we saw a significant decrease in locomotion activity in male mice compared to female mice. Acute stress had no effect on fear-related behavior or metabolic parameters. We also interestingly found differential abundance of MPPs and downstream GMPs and MDPs in the BM of HFD fed mice in a sex-specific manner. We also see sex-specific change in the number of immune cells including neutrophils and macrophages in the gWAT, aorta and heart. snRNAseq of the VMH of HFD mice revealed a cell type specific differences in female and male mice indicating the employment of differential pathways in the manifestation of fear-related responses, metabolic and immune dysfunction.

Our finding are in line with studies that have shown that stress and stress-associated disorders like PTSD can cause disruptions in the neuroendocrine [29–31], sympathetic [32–34], metabolic [35–37], and inflammatory systems [38–40], and lead to adverse changes in lifestyle that further contributes to deteriorating health [41–44]. Stress disrupts the established equilibrium, triggering physiological and metabolic adaptations in the body to address the demands challenging its homeostasis. Activating the hypothalamic-pituitary-adrenal (HPA) axis and the sympathetic nervous system (SNS) facilitates these adaptations. Subsequent exposure to repeated stressors following an initial extreme event can impose an excessive burden on these physiological systems, potentially resulting in various disease states as they struggle to cope with sustained stress. But chronic stress can create an undue burden on these systems to cope with the stressor leading to various disease states. The cascade of events stemming from this accumulated allostatic load contributes to the emergence of metabolic and inflammatory disorders [45]. Negative alterations in health can also be linked to unhealthy lifestyle choices, such as a high-fat diet, which may potentially stem from a fear of encountering environmental triggers that elicit symptoms of post-traumatic stress disorder (PTSD). This dietary preference might serve as a coping mechanism adopted by the individual [46]. Our knowledge of how diet and stress in conjuncture, affect behaviour, metabolism, and inflammation have been limited. Our finding that HFD and repeated stressor alters the number of immune cells are in line with alteration in immune modulating stress hormone cortisol [47], that in turn affect cognitive function [15–17, 48–50], and promote inflammation [50, 51].

We observed that male mice on the HFD displayed impaired fear memory extinction, as evidenced by their failure to exhibit a significant reduction in freezing behavior compared to the repeated shock group at week 14. This suggests that HFD inhibits fear extinction processes in male mice. This finding is consistent with previous research indicating a link between diet, particularly high-fat diets, and cognitive functions, extending it to fear memory extinction. Importantly, no such impairment was observed in females on the HFD, as they exhibited fear memory extinction similar to the chow-fed females. This could hint at diet being one of the key factors as to why PTSD develops in some but not in others. Many models developed from years of research have proposed that impaired extinction of the fear memory associated with the trauma is crucial for the development of PTSD symptoms and is directly correlated with symptom severity [52–54]. Diet could potentially be a risk factor for impaired extinction learning in PTSD and there exists a need to further investigate the link.

Our exploration into the effects of diet and stress on weight gain and blood glucose levels revealed sex-specific differences. While both male and female mice on the HFD exhibited increased weight gain over the 14-week period, males displayed a higher rate of weight gain compared to females, regardless of the stress condition. Additionally, blood glucose levels were influenced by both diet and sex. Unsurprisingly we see that HFD animals show higher blood glucose levels when injected with a bolus of glucose. However, HFD-fed females displayed elevated blood glucose levels compared to chow-fed females, particularly at earlier time points. Remarkably, in the absence of stressors, blood glucose levels returned to baseline chow-fed levels in HFD-fed females by week 14. Yet, HFD-repeated stressed females continued to show high levels of blood glucose, indicating glucose intolerance and insulin resistance phenotypes. Additionally, HFD-fed males continued to show elevated blood glucose levels even in the absence of stress and this difference was observed even in week 14. These results suggest a complex interplay between diet, sex, and stress in modulating metabolic outcomes. As mounting evidence indicate associations between PTSD and diabetes [55, 56], our results suggest a complex interplay between diet, sex, and stress in modulating metabolic outcomes. These disruptions lead to increased visceral adiposity, inflammation, and insulin resistance which might, in turn, perpetuate further irregularities in HPA axis functioning, creating a harmful cycle of deteriorating health [57].

In a state of homeostasis, food consumption usually aligns with energy expenditure. However, when rodents are exposed to a HFD or experience chronic stress, it can alter the central and peripheral mechanisms that regulate energy balance, leading to hyperphagia and greater accumulation of adipose tissue [58, 59]. However, there is a huge gap in knowledge on how exposure to both HFD and repeated stress could alter energy metabolism. In our studies we investigated the compounding effects of HFD and repeated stress in female and male mice by employing indirect calorimetric measures of various metabolic parameters. The overall daily energy expenditure can be broken down into various components, including basal or resting energy expenditure, physical activity, thermoregulation, and the thermic effect of food. Males on HFD show higher energy expenditure contrary to previous studies by Huang et al., [60] showing higher energy expenditure in HFD-fed female mice than male mice. This effect could be accounted for by the effect of both short-term stress induced by SEFL in NRS male mice on HFD and chronic stress effects in RS male mice on HFD. Implying an undue effect of stress, in any form, has a compounded effect with diet in disrupting energy homeostasis since we do not see these effects in chow-fed animals. This is also an extremely interesting finding given that we see a decrease in locomotion in these HFD-fed RS male mice. More robust studies are required to understand the increased energy expenditure seen without the physical demand of locomotion. Studies have also shown that RER decreases with HFD [61], but we see this effect come through only in female mice fed HFD emphasizing the need to rethink the importance of studying sex-differences. We also see a trend towards increased food consumption in chow-fed RS female mice recapturing clinical aspects of PTSD and eating disorder comorbidity [62, 63].

The findings from acute stress-induced experiments indicate that HFD-induced weight gain is associated with both sex and diet effects. While both male and female mice on HFD gained more weight than their chow-fed counterparts, males gained weight at a significantly higher rate. Moreover, fat mass in males on HFD was significantly higher compared to males on chow correlated with an increase in energy expenditure. These results suggest that the impact of HFD on weight gain and fat mass might be more pronounced in males. Interestingly, the analysis of freezing behavior did not show any sex or diet effect, indicating that acute stress might have a more consistent impact on freezing behavior regardless of sex or diet. This could indicate that the impact of acute and chronic stress on metabolic health differs significantly and HFD could exacerbate these effects.

We see a varying effect of sex on neutrophils and macrophages in various peripheral tissues. Nuetrophils in the gWAT of male mice fed a HFD may actively contribute to obesity-related inflammation. Neutrophils are innate immune cells that enter white adipose tissue, releasing inflammatory factors that can attract macrophages and other immune cells Neutrophils also play a key role in the development of vascular diseases aortic aneurysms [64, 65] and inflammation-induced atherosclerosis [66, 67]. Increased levels of neutrophils in the aorta of HFD-fed female mice could point us towards the manifestation of cardiovascular pathologies. While increased levels of macrophages in the heart tissue of HFD-fed male mice could indicate the initiation of heart inflammation [68]. While our studies have implicated a sex-specific manifestation of comorbid pathologies in HFD-fed mice, the delineation of effect of stress is lacking. Further studies are required to delve into the working mechanisms in the development of these various cardiac and metabolic diseases.

Hematopoietic stem cells (HSCs) in the BM produce erythroid, myeloid, and lymphoid cells and possess the ability to self-renew and differentiate into multiple cell types but are rare and mostly quiescent, dividing infrequently in the healthy state [69, 70]. Hematopoiesis is known to be impacted by HFD [71, 72] and various forms of stressors [73, 74]. The majority of continuous blood cell production is carried out by downstream multipotent progenitors (MPPs) which have more limited self-renewal capabilities and a more restricted lineage differentiation potential [75, 76]. MPPs sustain hematopoiesis by giving rise to common myeloid progenitors (CMPs), granulocyte–monocyte progenitors (GMPs) and myeloid dendritic progenitors (MDPs) [24, 77]. The reduction in myeloid-biased MPPs that can generate myeloid cells (i.e., MPP2 and MPP3), in HFD males and HFD-RS female mice could be accounted for by either an aberrant overproduction and rapid differentiation into downstream myeloid lineages cells. This theory fits well with the observed increase in the next hierarchical product of hematopoiesis the GMPs in HFD-fed male mice and the increase in MDPs in HFD-NRS female mice. Further studies are essential to tease apart the changes in hematopoietic continuum in response to diet and stress challenges.

Our snRNA-seq analysis of the VMH from male and female mice exposed to a high-fat diet with chronic restraint stress (RS-HFD) revealed the impact of both chronic stress and obesity on various cell types within the VMH, potentially influencing metabolic phenotypes. Data obtained from pooled VMH regions of male and female mice indicated differences in cell fractions among both neuronal and non-neuronal populations. While the overall qualitative clustering was similar between males and females, the analysis based on regulon activity and transcriptional signatures of VMH cell types unveiled an active transcriptional state in glial cells, including oligodendrocytes, astrocytes, microglia, and oligodendrocyte precursors. Male mice exhibited a higher astrocytic state with increased expression of *Gfap* and *Pomc*, while female mice showed a higher fraction of microglia and oligodendrocytes, suggesting a differentiation activation of glial cells in both sexes. Differential gene expression analysis revealed upregulation and downregulation of numerous genes in the glial cells of both males and females. Increased expression of the genes encoding orexin (*Hcrt*) in microglia and the gene encoding pro-Melanin concentrating hormone (*Pmch*) in oligodendrocytes was observed in females. Orexin and PMCH are neuropeptides that play major roles in wakefulness, and in addition to their actions in the central nervous system (CNS), they also regulate appetite, feeding, gastrointestinal mobility, energy balance, metabolism, blood pressure, neuroendocrine, and reproductive functions in various peripheral organs. Our findings indicated that RS-HFD females exhibited slightly increased locomotion and food intake but significantly decreased energy balance. Female mice under RS-HFD conditions showed marked activation of microglia. Moreover, our results demonstrated a significant glucose intolerance in female mice compared to those on a normal diet (NS-HFD) and reduced energy expenditure despite lower body weight. GWAS analysis revealed a higher trait of Alzheimer’s in female mice compared to male mice based on gene expression analysis. Microglia are implicated in diabetic and Alzheimer’s phenotypes, suggesting that chronic RS-HFD may contribute to a metabolic disorder in females. Similarly, our cell-cell communication analysis showed that female mice also exhibited stronger connections from microglia to glutamatergic neurons, oligodendrocyte precursors, and astrocytes.

Overall, our study has laid the groundwork for further comprehension of the mechanisms underlying metabolic health and the onset of stress-related disorders. Subsequent research is essential to explore the underlying mechanisms driving these sex-specific effects, encompassing neural, hormonal, and molecular factors, providing a clearer understanding of these complex interactions.

## Materials and Methods

### Animals

All experimental procedures were conducted in accordance with the guidelines set by the Institutional Animal Care and Use Committee at the Icahn School of Medicine at Mount Sinai. Ten-week-old strains of C57BL/6 mice, both males and females were ordered from Jackson Laboratories. Mice were housed in a temperature-controlled room under a 12-h light–12-h dark cycle and under pathogen-free conditions. Mice were fed a chow diet except when indicated that mice were placed on a high-fat diet (60% fat by calorie). At the time of euthanization, tissues, and blood were collected by cardiac puncture, frozen immediately, and stored at -80°C.

### Behavior Tests

#### Measure of Freezing and Shock Reactivity

Freezing is a complete lack of movement except for when the animal is breathing [22, 78] Freezing was measured using the EthoVision XT software that uses real-time video recording. The software measures freezing by a method called activity detection. The software detects changes at the pixel level from one video frame to the next. The threshold for activity detection is set at 0.02%. Freezing is counted when the activity score remains below this threshold for 1 sec. Percentage freezing = Freezing Time /Total Time×100 for the period of interest. Data are presented as mean percentages (+/− SEM). Shock reactivity is an index of pain reactivity to the incoming foot shock. Activity burst velocity is a reliable predictor of foot-shock intensity and may be a useful indicator of pain reactivity. We calculate shock reactivity by the velocity of movement during the 1-s period during the shock on Day 2. Velocity in inch/s (+/− SEM) was computed for the period of interest.

### Acute Stress

We ran a two-day acute stress model to induce acute stress in mice. Mice that were placed on either Chow or HFD either received acute stress (10 randomized foot-shocks over 1 hour) or no stress. On day 1, mice were placed in a fear conditioning chamber Context A. In context A, the mice were transported to the chambers directly from their home cages. Within the chamber, the conditional stimulus was presented in the form of white light, white noise, and the olfactory cue of a wet paper towel. The mice were placed in this chamber for a period of 1 hour, where the group of mice receiving acute stress was presented with 10 random foot shocks of 1mA delivered during a one-second period. The control animals for both dietary groups did not receive any foot shocks and hereby will be referred to as the no stress group. Fear generalization was tested on day 2 in context B. In this context, the mode of transport, light, visual, and olfactory cues was made drastically different. Mice were transported into the behavior room in individual opaque containers. An additional source of light was added on top of the chamber boxes, the walls of the chamber were changed to red, and the olfactory cue within the chamber now was that of 1% acetic acid. The grid floors are, however, kept the same from day 1 to day 2. The mice were placed in this new context for a period of 4 minutes where their fear was measured as an index of freezing.

### Stress-Enhanced Fear Learning

SEFL is a robust and powerful rodent model that recapitulates many clinical aspects of PTSD [22]. This model is carried out over a period of 3 days. On day 1, fear conditioning chambers are set up to serve as Context A. This context is followed exactly as described above in the acute stress condition. All mice are placed in the chamber for 1 hour and they are presented with 10 random foot shocks of 1 mA during this period. For days 2 and 3, context B is followed as described above. On day 2, the mice are placed in the chambers for a period of 4 minutes and 30 seconds. The mice receive 1-foot shock of 1 mA at the end of the 4th minute to serve as a reminder. On day 3, their fear is measured as an index of freezing in context B, by placing the mice in the chambers for a period of 8 minutes. Their percentage freezing is calculated from day 3. An additional measure of shock reactivity is calculated during the 4th-minute shock on day 2 using the same video-freeze system.

### Repeated Chronic Stress

After SEFL, both HFD and chow male and female animals were randomized to undergo repeated chronic stress or no repeated stress. The animals were placed week after week for 14 weeks in context B for 4 minutes. Repeated stress group received one foot shock at the end of 4^th^ minute while no repeated stress groups were allowed to extinguish the fear memory developed through SEFL.

### Open Field Light Gradient Anxiety Test

The mice were also tested on anxiety-like behavior in a modified open-field task that incorporated light gradient [79]. The open field light gradient test measured anxiety-like phenotype through parameters like distance traveled, velocity, and time spent in the dark zone. A rectangular, white, translucent polyethylene box (46 × 86 × 30 cm) was situated in a dark room. Three lamps are placed on one end of the box. A camera suspended above captures the animal’s activity. The test is divided into three phases during the 12-minute run. During the first four minutes, the lights are turned off and the animal can explore the open field in dark phase 1. During the next four minutes, the lamps are turned on and a light gradient is created in the open field. The zone closest to the light source called the “light zone” has an illumination index of ∼1200 lux. The “middle zone” has intermediate illumination of ∼50 lux and the zone farthest from the light source, the “dark zone” has an illumination index of ∼10 lux. In the last phase of the experiment, dark phase 2, the lights are turned off again. The locomotive activity, velocity, and time spent in the dark zone were calculated using the EthoVision XT software.

### Indirect Calorimetry using metabolic chamber

The mice were placed in an indirect calorimetry chamber for 72 hours for the purpose of studying the animal’s metabolic function after 10 weeks of diet/ diet+trauma administration in acute stress and repeated stress experiments respectively. Data collected from indirect calorimetry included oxygen consumption, carbon dioxide production, energy expenditure (EE), respiratory exchange ratio (RER), energy balance, food and water intake, locomotor activity, and body mass. For some indirect calorimetry measurements in the metabolic chambers such as oxygen consumption, carbon dioxide production, EE, and food and water intake, body weight was considered a covariate in statistical analysis. RER value and locomotion were analyzed without a covariate as they are known to be independent of body weight. This information highlights changes in metabolic physiology in animals placed in HFD/Chow after receiving either repeated trauma or acute stress. We analyzed the data with CalR [80] that considers activity, food intake, and other parameters allowing us to derive accurate indirect calorimetry values.

### Glucose Tolerance Test

For the glucose tolerance test, after 10 weeks and 14 weeks of feeding either HFD or Chow in both experimental groups, the mice fasted for 4 hours. The baseline blood glucose levels (mg/dL) in these mice were measured using one drop of blood collected from the tip of the tail and a glucometer. For the test, 1 g glucose/kg body weight of glucose was injected intraperitoneally (i.p.) into each mouse. Blood glucose levels were determined at times 0, 15, 30, 60, 90, and 120 min.

### Insulin Tolerance Test

For the insulin tolerance test (ITT), mice maintained on the HFD or Chow diet for 10 weeks and again after 14 weeks were fasted for 4 h before receiving i.p. injection of recombinant human insulin (1U/kg body weight). Blood glucose levels were determined at times 0, 15, 30, 60, and 90 min using a glucometer.

### Tissue Harvests

After completing all the above experiments, mice were food deprived for 4 hours, anesthetized with isoflurane, and rapidly decapitated. Multiple tissues were harvested and rapidly stored in a -80°C freezer. iWAT, gWAT, BAT, blood, liver, and brain were harvested from acute stress groups. In addition to these tissues, in repeated stress groups, the heart, aorta, and bone marrow were also harvested.

### Cell collection

Blood was collected and RBC lysis buffer (BioLegend) was used twice to lyse the red blood cells. After transcardiac perfusion with PBS (Thermo Fisher Scientific), organs (infarcted cardiac tissue and brain) were collected, minced and digested in a mixture of 450 U/mL collagenase I, 125 U/mL collagenase XI, 60 U/mL DNase and 60 U/mL hyaluronidase (Sigma-Aldrich) in PBS for 45 on a shaker (at 800 rpm) at 37 °C. Next, the digested organ was flushed through a 100 µm cell strainer. Brain suspensions were further purified using a Percoll density gradient (30% upper and 70% lower phase). The cell layer between the two phases was collected and washed in PBS. Bone marrow cells were flushed from the bone marrow cavities and brought into a single-cell suspension by pipetting up and down; then, RBC lysis buffer was used to lyse the red blood cells.

### Flow cytometry

Single-cell suspensions were stained in FACS buffer (0.5% BSA and 2 mM EDTA in PBS) containing fluorophore-coupled antibodies at a concentration of 1:700 at 4 °C for 30 min, unless otherwise indicated. To differentiate between live and dead cells, the cell suspensions were stained with Live/Dead Blue (ThermoFisher) at a concentration of 1:1,000 in PBS at 4 °C for 30 min. The following antibodies were used for flow cytometry: anti-CD45 (BioLegend, 30-F11), anti-CD11b (BioLegend, M1/70), anti-CD90.2 (Invitrogen, 53-2.1 and BioLegend, 30-H12), anti-B220 (BioLegend, RA3-6B2), anti-CD19 (BioLegend, 6D5), anti-CX3CR1 (BioLegend, SA011F11), anti-Ly-6G (BioLegend, 1A8), anti-Ly-6C (BioLegend, HK1.4), anti-f4/80 (BioLegend, BM8), anti-MHCII (BioLegend, M5/114.15.2), anti-CD64 (BioLegend, X54-5/7.1), anti-CD49b (BioLegend, DX5), anti-Ter119 (BioLegend, TER-119), anti-CD11c (BioLegend, N418), anti-CD127 (BioLegend, S18006K), anti-cKit (BioLegend, 2B8), anti-Sca-1 (BioLegend, E13-161.7), anti-CD135 (BioLegend, A2F10), anti-CD48 (BioLegend, HM48-1), anti-CD150 (BioLegend, TC15-12F12.2), anti-CD34 (eBioscience, RAM34), anti-CD16/32 (BioLegend, 93), anti-CD115 (BioLegend, AFS98), anti-BrdU (eBioscience, BU20A). Anti-Feeder Cells (Miltenyi Biotec, mEF-SK4), anti-podoplanin/gp38 (BioLegend, 8.1.1), anti-CD31 (BioLegend, 390). Mature cells were identified as: (1) Ly6CHi monocytes (CD45+ CD11b+ CX3CR1+ f4/80-Ly6CHi), (2) Neutrophils (CD45+ CD11b+ CX3CR1-Ly6G), (3) macrophages (CD45+ CD11b+ CX3CR1+ f4/80+ CD64+), (4) fibroblasts (CD45-CD31-gp38+ mEFSK4+), (5) T Cells (CD45+CD11b-CD90.2+), (6) B Cells (CD45+CD11b-B220+), (7) microglia (CD45midCD11b+). Progenitor cells were identified as: (1) LSK (CD45+Lin-Sca1+cKit+), (2) granulocyte-macrophage progenitor (CD45+Lin−cKit+Sca1−CD34+CD16/32HiCD115−), (3) monocyte-dendritic cell progenitor (CD45+Lin−cKit+Sca1−CD34+CD16/32HiCD115+), (4) common myeloid progenitor (CD45+Lin−cKit+Sca1−CD34-CD16/32mid). Lineage was identified as; Lin: B220, CD19, CD49b,Ter119, CD90.2, CD11b, CD11c, Ly6G, CD127. Data were acquired using a Cytek Aurora (Cytek) and analyzed with FlowJo (Tree Star).

### RNA purification, cDNA synthesis, and RT-qPCR

RNA was isolated from brown adipose tissue (BAT) and liver using phenol-chloroform extraction. After isolated, the RNA pellet was washed and resuspended in diethylpyrocarbonate (DEPC) water at a concentration of 200 ng/μL. RNA samples were reversely transcribed to cDNA using a high-capacity cDNA reverse transcription kit (Applied Biosystems). Real Time qPCR was performed by using a real-time PCR SYBR green master mix (Diagenode). Samples were run and analyzed on a Quantstudio 5 (Applied Biosystems). The qPCR targets were normalized to the expression of the housekeeping gene 36B4.

### Single-nuclei RNA sequencing (snRNA-seq)

Starting with the fresh-frozen VMH, we obtained approximately 100 mg of tissue by cutting brain punches, and the VMH regions from five mice were combined. Single nucleus gene expression sequencing was performed on the samples using the Chromium platform (10x Genomics, Pleasanton, CA) with the Next GEM Single cell 3’GEX Reagent kit. Briefly, nuclei were isolated from frozen tissue using the Singulator instrument (S2 Genomics, Livermore, CA) using the “Low Input Nuclei Isolation” protocol, where Mixing was modified to “Top” type and “Slow” speed. Additionally, disruption was performed via the “Dounce” method at “Medium” speed. Once run, the 3-4 mL of recovered supernatant was centrifuged at 300 xg for 4 min at 4oC and resuspended in 100uL PBS + 0.04% BSA, followed by filtration using a 40um cell strainer. Following loading of ∼10,000 healthy nuclei, Gel-Bead in Emulsions (GEMs) were generated on the sample chip in the Chromium controller. Barcoded cDNA was extracted from the GEMs by Post-GEM RT-cleanup and amplified for 12 cycles. Amplified cDNA was fragmented and subjected to end-repair, poly A-tailing, adapter ligation, and 10X-specific sample indexing following the manufacturer’s protocol. Libraries were quantified using Bioanalyzer (Agilent) and QuBit (Thermofisher) analysis. Libraries were sequenced using a 2x100PE configuration on a NovaSeq instrument (Illumina, San Diego, CA) targeting a depth of 50,000-100,000 reads per nucleus. Sequencing data was aligned and quantified using the Cell Ranger Single-Cell Software Suite (version 7.1.0, 10x Genomics) against the provided mm10 reference genome using default parameters, including introns.

### snRNA-seq pre-processing and quality control

To obtain digital gene expression matrices (DGEs) in sparse matrix representation, paired end reads from the Illumina NOVA seq were processed and mapped to the mm10 mouse genome using 10X Genomics’ Cell Ranger v3.0.2 software suite. Briefly, .bcl files from the Mount Sinai sequencing core were demultiplexed and converted to fastq format using the ‘mkfastq’ function from Cell Ranger. Next, the Cell Ranger ‘counts’ function mapped reads from fastq files to the mm10 reference genome and tagged mapped reads as either exonic, intronic, or intergenic. Only reads which aligned to exonic regions were used in the resulting DGEs. After combining all four sample DGEs into a single study DGE, we filtered out cells with (1) UMI counts < 700 or > 30,000, (2) gene counts < 200 or > 8,000, and (3) mitochondrial gene ratio > 10%. This filtering resulted in a dataset consisting with approximately 2,300 – 4,650 cells from each sample. A median of 2,411 genes and 7,252 transcripts were detected per cell.

### Identification of cell clusters

To achieve high resolution cell type identification and increased confidence in our cell type clustering we brought in external publicly available data. The single cell expression profiles were projected into two dimensions using UMAP method [81] for community detection was used to assign clusters. This integrated data was only used to identify and define the cell types. All plots which are not explicitly designated as integrated with at least one external dataset and all downstream analyses (e.g. differential expression analyses) were conducted on non-integrated data to retain the biological effect of the cold treatment. Visualization of the non-integrated data was conducted on a subsampled dataset where all samples had the same number of cells to give an equal weight to each sample, however, all downstream analyses (e.g. differential expression analyses) leveraged the full dataset.

### Cell type-specific marker gene signatures

Cell type-specific marker gene signatures were generated by identifying genes with expression levels two-fold greater (adjusted p-values < 0.05) than all other cell types. To ensure consistency across samples, Seurat’s FindConservedMarkers function (Wilcoxon rank sum test with a meta p-value) was applied across each sample.

### Resolving cell identities of the cell clusters

To identify the cell type identity of each cluster, we used a curated set of canonical marker genes derived from the literature to find distinct expression patterns in the cell clusters. Clusters which uniquely expressed known marker genes were used as evidence to identify that cell type. Cell subtypes which did not express previously established markers were labeled by both general cell type markers and novel markers obtained with Seurat’s FindConservedMarkers function were used to define the cell subtype.

### Differential gene expression analysis

Within each identified cell type or subtype, cold treated and room temperature single cells were compared for differential gene expression using Seurat’s FindMarkers function (Wilcoxon rank sum test) in a manner similar to Li et al. [82]. DEGs were identified using two criteria: (i) an expression difference of >= 1.5-fold and adjusted p-value < 0.05 in a grouped analysis between room temperature mice (n = 2) and cold treated mice (n = 2); (ii) an expression difference of >=1.25 fold and consistent fold change direction in all 4 possible pairwise combinations of cold-treated vs room temperature mice.

### Gene regulatory network inference

Gene regulatory network inference was performed with pySCENIC [83] following the workflow described by Van de Sande et al [84]. Briefly, starting with counts data, gene modules which are co-expressed with transcription factors were identified with GRNBoost2 [85]. Next, candidate regulons were created from transcription factor – target gene interactions and indirect targets were pruned based on motif discovery with cisTarget [83]. Finally, regulon activity was quantified at cellular resolution with AUCell [83] which allowed for the prioritization of regulons for each cell type based on the quantified activity.

### Statistical Analysis

Data are shown as mean±S.E.M. Distribution was assessed by Shapiro-Wilk test. Significance was determined by a two-tailed unpaired t test (parametric distribution), 2way ANOVA Bonferroni *post hoc* test, ANCOVA, or by a Mann-Whitney test (non-parametric distribution). Significance was set at an alpha level of 0.05.

## Acknowledgements

P.R is supported by R00DK114571, NIDDK-supported Einstein-Sinai Diabetes Research Center (DRC) Pilot & Feasibility Award, and Diabetes Action Research and Education Foundation (DAREF) Grant # 501 (PR). scRNA-seq library preparation and sequencing were performed under the the Illumina-Sinai Award (PR). A.K.R is supported by R21DK129908. The funders had no role in study design, data collection and interpretation, or the decision to submit the work for publication.

