## Supplemental Information for "Sexually dimorphic role of diet and stress on behavior, energy metabolism, and the ventromedial hypothalamus"

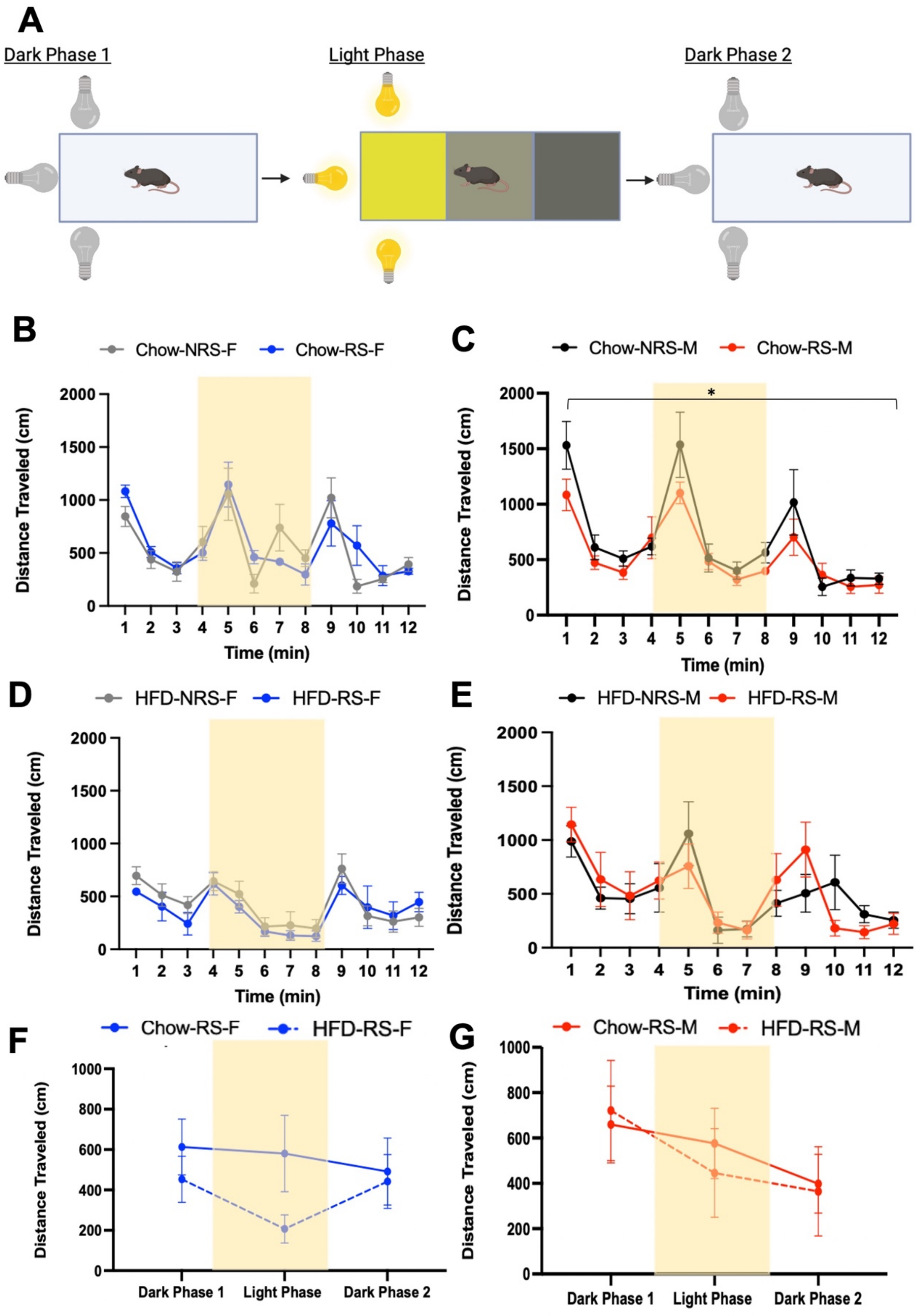

**Supplemental Fig 1. Repeated Shock and HFD induces an anxiety-like phenotype in female mice. A.** Experimental design of open field light-gradient task. **B.** Distance travelled over the 12 minute in chow fed female mice (N=5, Two way ANOVA, $F_{1,96} = 0.128$, $p>0.05$). **C.** Distance travelled over the 12 minute in chow fed male mice (N=5, Two way ANOVA, $F_{1,96} = 6.853$, $p<0.05$)**. D.** Distance travelled over the 12 minute in HFD fed female mice (N=5, Two way ANOVA, $F_{1,96} = 1.811$, $p>0.05$). **E.** Distance travelled over the 12 minute in HFD fed male mice (N=4/5, Two way ANOVA, $F_{1,84} = 0.044$, $p>0.05$). **F.** Average distance travelled during the three phases of the behavioral task in HFD/Chow fed RS female mice (N=5, Two way ANOVA, $F_{1,24} = 2.88$, $p=0.09$). **G.** Average distance travelled during the three phases of the behavioral task in HFD/Chow fed RS male mice (N=5, Two way ANOVA, $F_{1,24} = 0.054$, $p>0.05$).

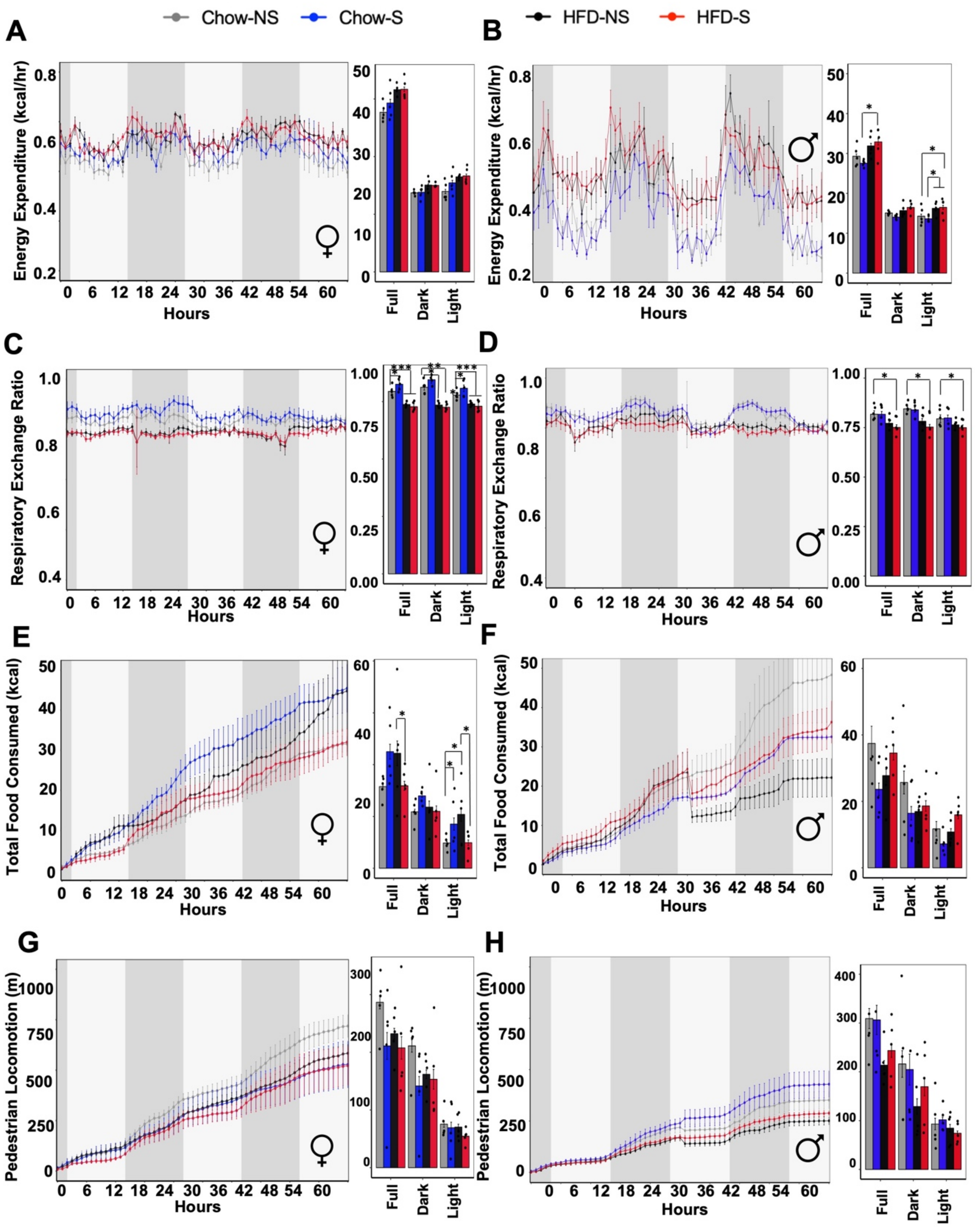


**Supplemental Fig. 2. Sex and acute stress effects on mice that are fed chow/HFD. A.** Energy expenditure (EE) (kCal/hr) of HFD/chow fed female mice during 60 hours of metabolic chamber housing (N=5. One way ANOVA, p>0.05). **B.** Energy expenditure (EE) (kCal/hr) of HFD/chow fed male mice during 60 hours of metabolic chamber housing (N=5, One way ANOVA, *p<0.05) (*Post hoc* comparison, N=4,5, Tukey's multiple comparison test, *p<0.05).

**C.** Respiratory exchange ratio (RER) in HFD/chow fed female mice (N=5, One way ANOVA, ***$p<0.001$) (*Post hoc* comparison, N=5, Tukey's multiple comparison test, *$p<0.05$, ***$p<0.001$). **D.** RER in males on HFD/Chow diet (N=4,5, One way ANOVA, ***$p<0.001$) (*Post hoc* comparison, N=5, Tukey's multiple comparison test, *$p<0.05$). **E.** Total food consumption (kCal) in HFD/chow fed female mice (N=5, one way ANOVA, ***$p<0.001$) (*Post hoc* comparison, N=5, Tukey's multiple comparison test, *$p<0.05$, **$p<0.01$, ***$p<0.001$). **F.** Total food consumption (kCal) in HFD/chow fed male mice (N=5, One way ANOVA, $p>0.05$). **G.** Pedestrian locomotion (m) in HFD/chow fed female mice (N=5, One way ANOVA, $p>0.05$). **H.** Pedestrian locomotion (m) in HFD/chow fed male mice (N=5, One way ANOVA, $p>0.05$).

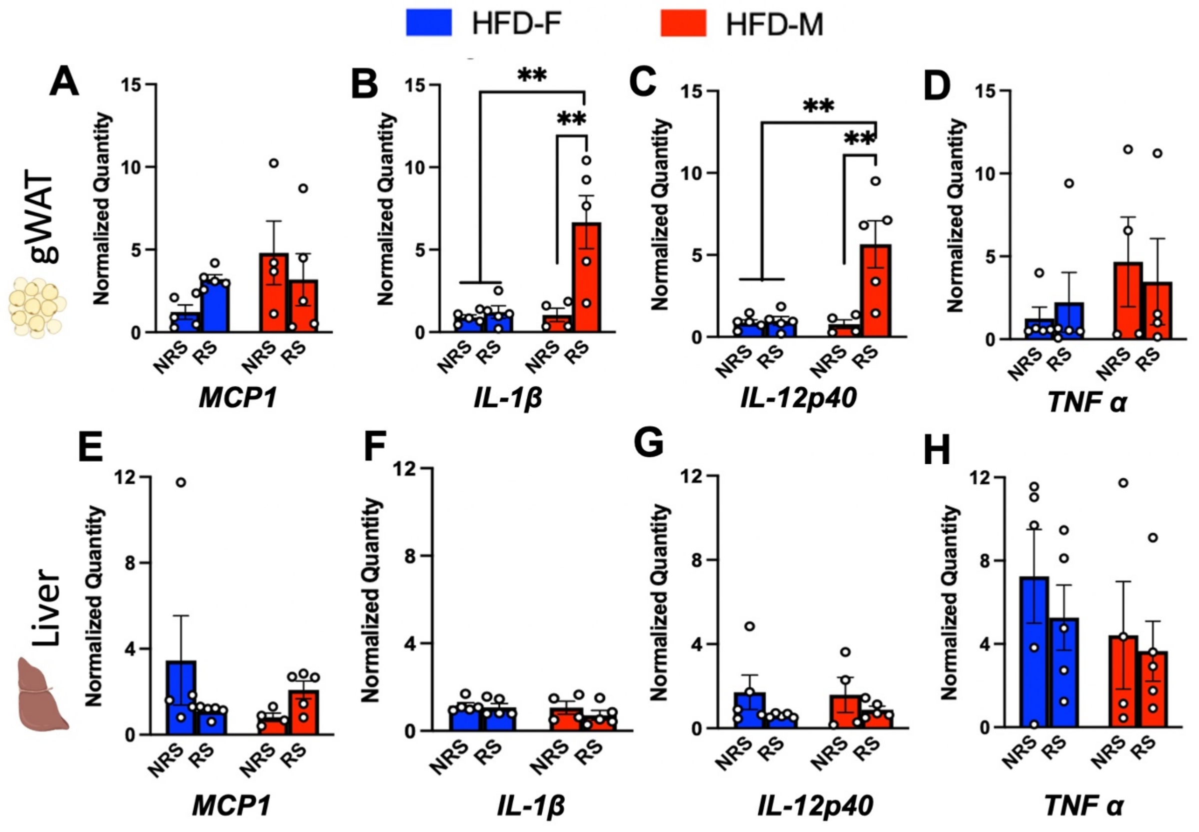


**Supplemental Fig 3. HFD induces upregulation of inflammatory markers in the peripheral tissue. A.** Normalized quantification of MCP-1 gene expression in gWAT of HFD fed animals (N=9, two way ANOVA, $F_{1,15} = 2.248$, $p>0.05$) **B.** Quantification of pro-inflammatory cytokine IL-1β in gWAT of HFD mice (N=9, two way ANOVA, $F_{1,15} = 9.796$ &11.08 respectively, **$p<0.01$) (*Post hoc* comparison, N=4/5, Tukey's multiple comparison test, **$p<0.01$). **C.** Quantification of cytokine IL-12p40 in gWAT in HFD fed mice (N=9,10, two way ANOVA, $F_{1,15} = 9.796$ & 11.08 respectively, *$p<0.05$ & **$p<0.01$ respectively) (*Post hoc* comparison, N=4/5, Tukey's multiple comparison test, **$p<0.01$). **D.** Quantification of TNF 〈 levels in HFD fed animals in the gWAT(N=9, two way ANOVA, $p<0.05$). **E,F,G &H.** Quantification of MCP-1, IL-1β, IL-12p40 or TNF 〈 gene expression levels in liver of HFD-fed animals (N=9, two way ANOVA, $p<0.05$).

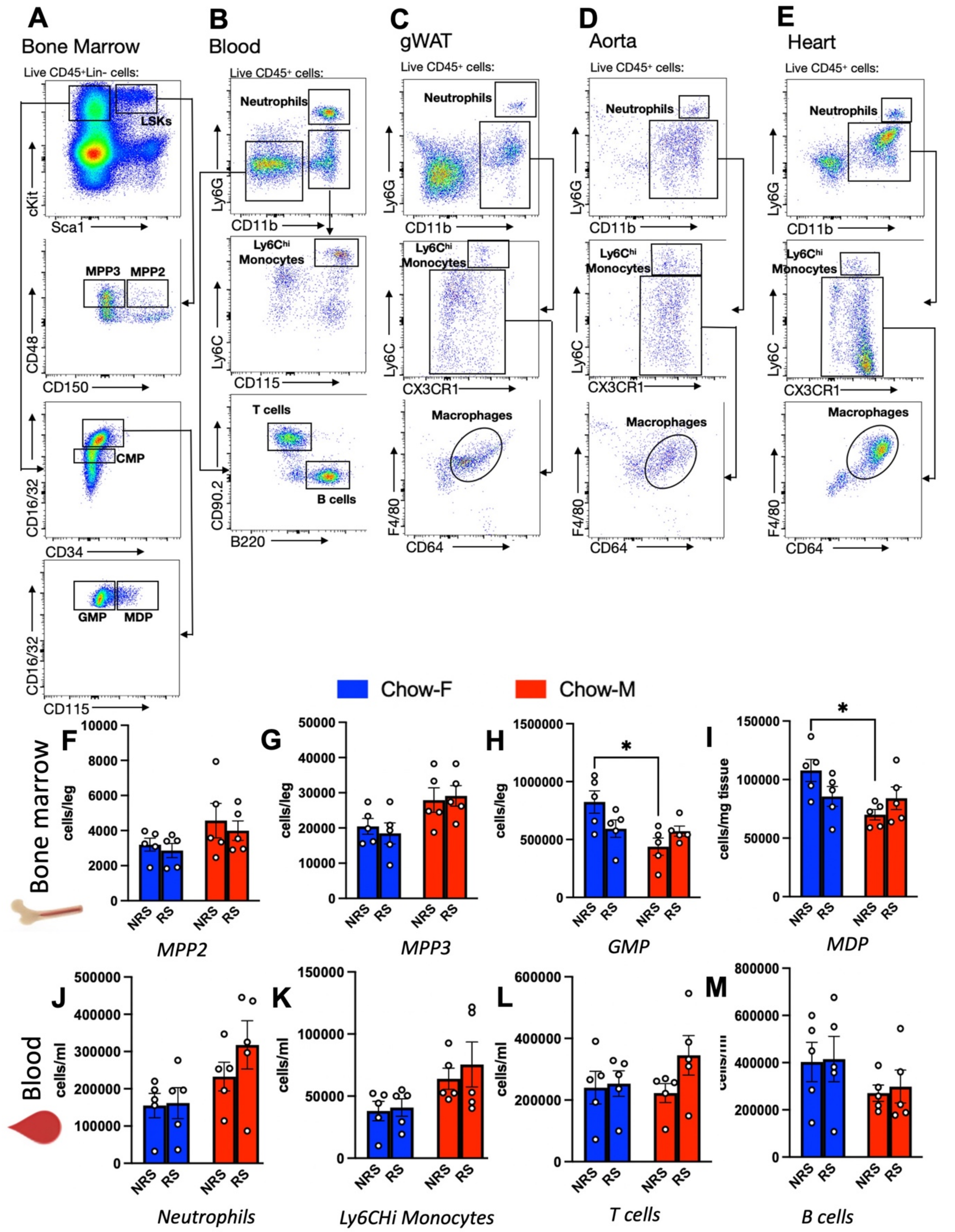


**Supplemental Fig. 4 Chow diet has minimal effects on peripheral myeloid lineage cells and inflammatory markers. A-E.** Gating strategies for flow cytometric data analysis. **F.** FACS analysis and further quantification of MPP2 cells in the BM of HFD (N=10, two way

ANOVA, $F_{1,16}$ = 4.007, $p>0.05$). **G.** Quantification of MPP3 cells in the BM of chow-fed animals (N=10, two way ANOVA, $F_{1,16}$ = 9.504, $p>0.05$) **H.** Quantification of GMPs in BM of chow fed animals (N=10, two way ANOVA, $F_{1,16}$ = 7.478, *$p<0.05$) (*Post hoc* comparison, N=5, Tukey's multiple comparison test,* $p<0.05$). **I.** Quantification of MDPs in the BM of chow animals (N=10, two way ANOVA, $F_{1,16}$ = 5.604, *$p<0.05$) (*Post hoc* comparison, N=5, Tukey's multiple comparison test,* $p<0.05$). **J.** Quantification of neutrophil levels in the blood of chow fed mice (N=9, two way ANOVA, $F_{1,16}$ = 6.46, *$p<0.05$ (*Post hoc* comparison, N=5, Tukey's multiple comparison test, $p>0.05$). **K.** Quantification of Ly6CHi monocytes in the blood of chow fed mice (N=10, two way ANOVA, $F_{1,16}$ = 6.46, *$p<0.05$) (*Post hoc* comparison, N=5, Tukey's multiple comparison test, $p>0.05$). **L.** Quantification of T cells in the blood of chow-fed animals (N=9, two way ANOVA, $p>0.05$). **M.** Quantification of B cells in the blood of chow-fed animals (N=10, two way ANOVA, $p>0.05$).

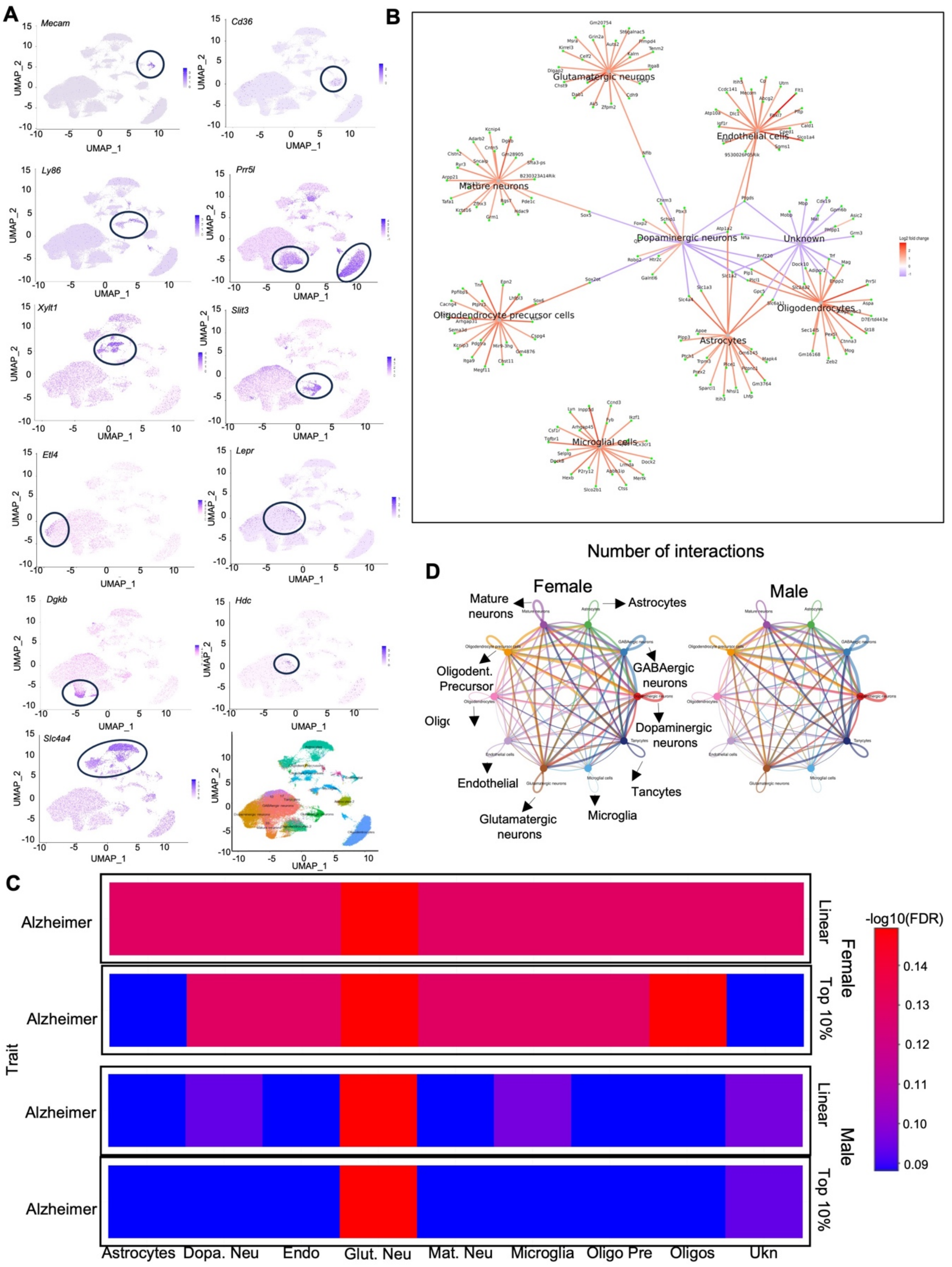


**Supplemental Fig. 5. scRNA-seq from VMH show celltype specific gene expression, cellular interaction and GWAS phenotype in males and females. A.** The UMAP plot depicts the distribution of annotated clusters from the combined dataset based on indicated marker genes. **B.** Analysis of cell-type-specific feature genes and shared gene connections within labeled clusters. **C.** A GWAS study reveals Alzheimer traits associated with differentially expressed genes (DEGs) in male and female mice across specified cell types. **D.** The interactome illustrates the number of interactions between cell types, derived from ligand-receptor interactions in males and females.
